# Conserved amino acid residues and gene expression patterns associated with the substrate preferences of the competing enzymes FLS and DFR

**DOI:** 10.1101/2023.11.05.565693

**Authors:** Nancy Choudhary, Boas Pucker

## Abstract

**Background:** Flavonoids, an important class of specialized metabolites, are synthesized from phenylalanine and present in almost all plant species. Different branches of flavonoid biosynthesis lead to products like flavones, flavonols, anthocyanins, and proanthocyanidins. Dihydroflavonols form the branching point towards the production of non-colored flavonols via flavonol synthase (FLS) and colored anthocyanins via dihydroflavonol 4-reductase (DFR). Despite the wealth of publicly accessible data, there remains a gap in understanding the mechanisms that mitigate competition between FLS and DFR for the shared substrate, dihydroflavonols.

**Results:** An angiosperm-wide comparison of FLS and DFR sequences revealed the amino acids at positions associated with the substrate specificity in both enzymes. A global analysis of the phylogenetic distribution of these amino acid residues revealed that monocots generally possess FLS with Y132 (FLS_Y_) and DFR with N133 (DFR_N_). In contrast, dicots generally possess FLS_H_ and DFR_N_, DFR_D_, and DFR_A_. DFR_A_, which restricts substrate preference to dihydrokaempferol, previously believed to be unique to strawberry species, is found to be more widespread in angiosperms and has evolved independently multiple times. Generally, angiosperm FLS appears to prefer dihydrokaempferol, whereas DFR appears to favor dihydroquercetin or dihydromyricetin. Moreover, in the FLS-DFR competition, the dominance of one over the other is observed, with typically only one gene being expressed at any given time.

**Conclusion:** This study illustrates how almost mutually exclusive gene expression and substrate-preference determining residues could mitigate competition between FLS and DFR, delineates the evolution of these enzymes, and provides insights into mechanisms directing the metabolic flux of the flavonoid biosynthesis, with potential implications for ornamental plants and molecular breeding strategies.

## Introduction

Flavonoids comprise a range of different subclasses including flavones, flavonols, anthocyanins, and proanthocyanidins, all of which have their own biological functions [1]. Flavonoids exhibit a remarkable diversity, encompassing a vast array of compounds that possess distinct properties, conferring specific evolutionary benefits [2]. The biosynthesis of these compounds involves a complex network of enzymatic reactions. Mutants with observable phenotypic variations in flavonoid biosynthesis have served as valuable model systems for studying the intricacies of biosynthetic pathways and their regulatory mechanisms [3–6].

Anthocyanins are renowned for their role as red, purple, or blue pigments, contributing to the vibrant colors seen in flowers and fruits [1]. These color patterns serve to attract animals for important biological processes such as pollination and seed dispersal [7]. Flavonols have been described as co-pigments that contribute to the overall coloration of various plant structures [8,9]. On their own, flavonols are colorless or pale yellow and have been described as important factors in the UV stress response [10]. Additionally, flavonoid biosynthesis produces other compounds such as flavones and proanthocyanidins [11,12]. Flavones play a crucial role in signaling and pathogen defense [11], whereas proanthocyanidins are key contributors to seed coat pigmentation [13]. Beyond their pigmentation functions, flavonoids have diverse roles in response to environmental factors like drought [14], cold [15–17], and high light intensities [18,19], playing a vital physiological role in protecting against UV-induced oxidative stress [20,21], detoxifying reactive oxygen species (ROS) [22,23] and toxic metals [24], defending against pathogens [25], regulating symbiotic associations [26,27], and influencing auxin flux [28,29]. A theory says that the cytochrome P450 enzymes (like C4H, FNSII, F3’H, and F3’5’H) are anchored to the endoplasmic reticulum membrane and associate with other soluble enzymes in the cellular environment forming a “metabolon” [30]. Supported by biochemical experiments, flavonoid metabolons have been reported in a few plant species including soybean [31], Arabidopsis [32], kumquat[33], snapdragon, and torenia [34].

A range of different colors is provided by three classes of anthocyanins: pelargonidin (orange to bright red), cyanidin (red to pink), and delphinidin (purple or blue) glycosides [1]. These three classes are characterized by different numbers of hydroxyl groups at the B phenyl ring: pelargonidin (one), cyanidin (two), and delphinidin (three) [35]. The biosynthesis of anthocyanins involves the enzymatic conversion of colorless dihydroflavonols into leucoanthocyanidins by DFR [36]. ANS further converts leucoanthocyanidins into anthocyanidins. Recent research suggests the involvement of an anthocyanin-related glutathione S-transferase (arGST) in anthocyanin biosynthesis [37,38]. Anthocyanidins are transformed into anthocyanins through the addition of a sugar moiety by glycosyl transferases [39]. Subsequent modifications such as the addition of other sugar moieties [40] or acyl groups [41] are catalyzed by a large number of often promiscuous enzymes.

Dihydroflavonols are not exclusively channeled into anthocyanin biosynthesis but are also the substrate of the flavonol synthase (FLS) that produces kaempferol, quercetin, or myricetin depending on the available dihydroflavonol [42]. This leads to a competition for the dihydroflavonol substrates between DFR and FLS (**Fig 1**). Previous studies suggest that the ratio of FLS to DFR activity is an important factor in color determination [42–46]. A study across multiple flowering plants discovered that high FLS activity can result in a lack of anthocyanin pigmentation despite a functional anthocyanin biosynthesis machinery being present [45]. The flavonoid 3’-hydroxylase (F3’H) converts dihydrokaempferol into dihydroquercetin by adding one hydroxyl group in the C3’ position of the B ring, while the flavonoid 3’,5’-hydroxylase (F3’5’H) converts dihydrokaempferol into dihydromyricetin by adding hydroxyl groups at the C3’ and C5’ positions [47]. Differences in the stability of the different dihydroflavonols pose a challenge for quantitative *in vitro* studies [48].

**Fig 1:**
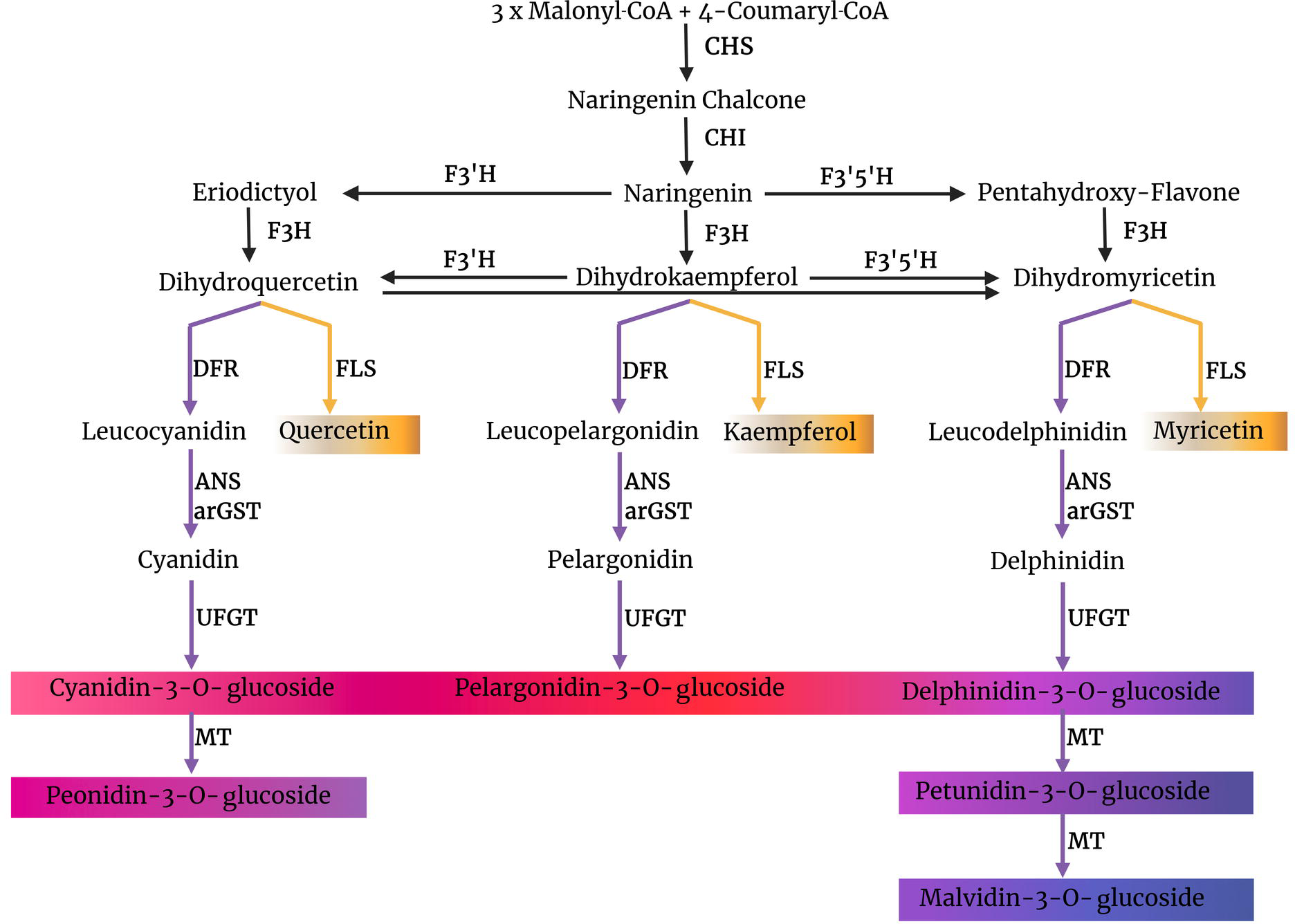
Simplified illustration of the flavonoid biosynthesis in plants showing the competition between FLS and DFR for common substrate, dihydroflavonols. CHS, chalcone synthase; CHI, chalcone isomerase; F3H, flavanone 3-hydroxylase; F3_′_H, flavonoid 3_′_-hydroxylase; F3_′_5_′_H, flavonoid 3_′_,5_′_-hydroxylase; FLS, flavonol synthase; DFR, dihydroflavonol 4-reductase; ANS, anthocyanidin synthase; arGST, anthocyanin-related glutathione S-transferase; UFGT, UDP-glucose flavonoid 3-O-glucosyl transferase; MT, methyltransferase.

The activity of different branches of flavonoid biosynthesis is controlled by transcriptional activation of the involved genes. *FLS* gene expression is activated by MYB transcription factors of the subgroup7 and this mechanism appears to be generally conserved across plant species with minor lineage-specific variations [49,50]. Recently, a pollen-specific activation of *FLS* by sugroup19 MYBs was described [51,52]. Transcriptional activation of DFR is controlled by an MBW transcription factor complex that is named after the three components: MYB, bHLH, and WD40 [53]. While research on *A. thaliana* mutants suggests that positive regulation dominates the gene expression control of flavonoid biosynthesis genes, several negative regulators have been identified through research on many different plant species [54].

Apart from the overall activity of FLS and DFR, substrate selectivity plays a crucial role in determining the production of flavonols and anthocyanins [55,56]. This selectivity is evident in species like *Freesia hybrida*, in which the presence of abundant kaempferol glycosides and delphinidin derivatives [57] indicates the separation of flavonol and anthocyanin biosynthesis based on the contrasting substrate preferences of DFR and FLS [57]. Despite testing various DFR copies in *Freesia hybrida*, none of them exhibited a preference for DHK, while all favored DHM as the substrate [57]. Some DFRs have been found to accept all three potential dihydroflavonols, indicating that the availability of specific substrates determines the class of anthocyanins produced. Other DFRs are incapable of accepting dihydrokaempferol as a substrate, resulting in the absence of orange anthocyanin-based pigmentation in species like petunia, apple, pear, hawthorn, rose, and cranberry [48,56,58]. The substrate specificity determining region was predicted by a multiple alignment of polypeptide sequences [59]. This region was further investigated based on the *Gerbera hybrida* DFR which can accept DHK and leads to orange flower pigmentation [60]. Two important amino acid residues implicated in the substrate specificity of DFR were identified based on chimeric *Gerbera hybrida* and *Petunia hybrida* DFRs followed by site-directed mutation experiments [56]. The relevance of all positions in a region of 26 amino acid residues was evaluated by permutation experiments. The mutation E145L of the *G. hybrida* DFR resulted in a non-functional enzyme while the N134L mutation restricted the substrate preference to dihydrokaempferol and excluded dihydroquercetin or dihydromyrecitin [56]. Many following studies confirmed the particular relevance of this N134 residue (position 133 in *Arabidopsis thaliana* DFR) and identified additional amino acid residues with specific properties [55,61–63]. In summary, N allows the acceptance of all dihydroflavonols, L restricts the substrate to DHK, D enables acceptance of DHQ and DHM, and A leads to a high DHK affinity (**Fig 2**).

**Fig 2:**
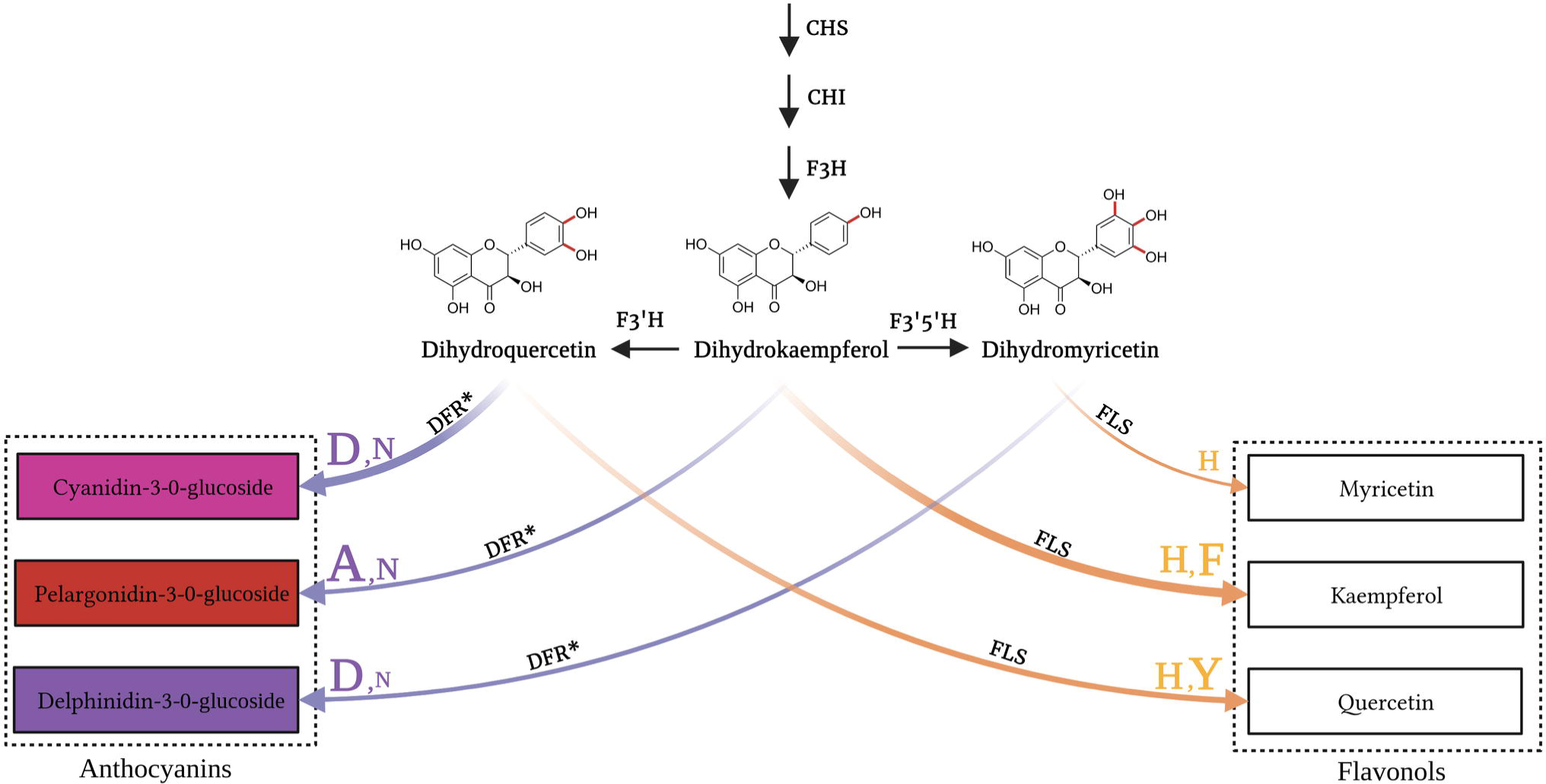
Illustration of the branching point in the flavonoid biosynthesis leading to anthocyanins and flavonols, respectively. Asterisk (*) shows that the reactions from dihydroflavonols to anthocyanins also required the additional enzymes anthocyanidin synthase (ANS), anthocyanin-related glutathione S-transferase (arGST) and UDP-dependent anthocyanidin 3-O-glucosyltransferase (UGT) downstream of DFR. DFR and FLS can process substrates with different hydroxylation patterns leading to three distinct products each. The characters at the arrows represent the previously reported substrate-preference-determining residues of FLS and DFR. The DFR residues, represented as purple characters depict the 133 position while the FLS residues, represented as yellow characters depict the 132 position in FLS (position according to *AtDFR* and *AtFLS1*).

While the substrate preference of DFR received substantial attention, little is known about the substrate preferences of FLS. Five specific residues, namely H132, F134, K202, F293, and E295, have been reported to be involved in the binding of the DHQ substrate [64]. Previous research in *A. thaliana* studied the impact of amino acid substitutions in AtFLS through incubation with DHQ. The mutants, H132F and H132Y exhibited 124% and 83% activity, respectively, compared to the wild-type (WT) [64]. The K_M_ values of mutants H132F and H132Y displayed an approximately 2-fold decrease compared to the WT counterpart [64]. Conversely, mutations in the other four residues led to a significant decrease in FLS activity. Interestingly, the four highly conserved residues were found to be crucial for substrate binding and catalytic activity. The H132 residue was found to be variable across different flavonol synthases, with some having F residues and others having H residues. These findings suggested that the H132 position may play a role in determining substrate preferences among different flavonol synthases [64]. It is postulated that the position of H132 interacts via a hydrogen bond with the B-ring hydroxyl group of the dihydroflavonol substrates [65]. Given the previous reports about the relevance of N133 in DFR and H132 in FLS, we will refer to them as substrate-preference determining residues throughout this manuscript, while not ruling out the contribution of additional amino acid residues.

While the model organism *A. thaliana* has only a single DFR gene [66], multiple DFR copies have been reported in many other plant species. For example, studies in buckwheat [67] and strawberry [68] have identified DFR copies that show substantial differences in their substrate preference and consequently also in the produced anthocyanin classes. Genetic analysis of *DFR1* and *DFR2* in buckwheat suggested that both genes are genetically linked, but the genetic distance does not suggest a recent tandem duplication [67]. Buckwheat *DFR2* differs from *DFR1* by an asparagine to valine substitution at a position that corresponds to 134 in the *Gerbera hybrida* sequence and the presence of an additional glycine between positions 138 and 139. Enzyme assays suggest that *DFR1* prefers DHK and DHM, while *DFR2* prefers DHQ. *DFR1* is expressed in most tissues, while *DFR2* expression is restricted to seeds and roots. Strawberry DFR1 has a high preference for DHK and does not accept DHQ and DHM (‘DHK specialist’), while DFR2 does not accept DHK as a substrate, but works on DHQ and DHM (‘DHK rejector’) [68].

The objective of this study was to reveal mechanisms that control and mitigate the substrate competition of FLS and DFR. A systematic analysis of substrate-preference-determining amino acid residues in the polypeptide sequences of FLS and DFR revealed differences between monocots and dicots. Cross-species gene expression revealed an almost mutually exclusive activity of *FLS* and *DFR*, which could be another mitigation mechanism.

## Methods

### Analyzed species and data sets

The coding sequences for 211 plant species were retrieved from Phytozome [69] and NCBI/GenBank [70] (Additional File 1). The selection of plant species encompassed a diverse taxonomic range, including 20 Thallophyta (algae and fungi), 4 Bryophyta (mosses), 1 Pteridophyta (ferns and fern allies), 11 Gymnosperms (naked seed plants), and 175 Angiosperms (flowering plants). A Python script [71] was used to translate the coding sequences into corresponding polypeptide sequences.

### Flavonoid biosynthesis gene identification

The polypeptide sequences were analyzed with KIPEs3 v0.35 [72,73] to identify flavonoid biosynthesis genes in each plant species. KIPEs3 itself supplied the bait sequences and essential residue information required for the analysis (flavonoid biosynthesis dataset v3); leading to the identification of candidate genes for *FLS* (*flavonol synthase*) and *DFR* (*dihydroflavonol 4-reductase*). In addition, flavanone 3-hydroxylase (*F3H)*, flavonoid 3’-hydroxylase (*F3’H)*, and flavonoid 3’5’-hydroxylase (*F3’5’H)* genes were identified in the investigated plant species. KIPEs3 also enabled the identification of *FLS*-like and *DFR*-like genes within the datasets.

### Multiple sequence alignment and phylogenetic tree construction

The DFR and DFR-like polypeptide sequences and the FLS and FLS-like polypeptide sequences were aligned separately using MAFFT v7.310 [74] with default penalties for gaps and the protein weight matrix of BLOSUM62 [75]. The alignments were generated using the L-INS-i accuracy-oriented method and incorporated local pairwise alignment. The amino acid alignments were translated back to codon alignments using pxaa2cdn [76]. The alignments were cleaned using pxclsq [76] to remove alignment columns with very low occupancy (<0.1). Multiple ANR and CCR sequences retrieved from GenBank were used as an outgroup for DFR tree construction. Previously characterized DFRs were also included. Multiple F3H and ANS sequences obtained from GenBank were used as an outgroup for FLS tree construction to facilitate a comprehensive investigation in the context of the 2-oxoglutarate-dependent dioxygenase (2-ODD) family. Previously described FLS sequences were also included. The best model of evolution was inferred using ModelFinder [77]. Maximum likelihood trees were constructed for deciphering *DFR* and *FLS* evolution using IQ-TREE v2.0.3 [78] with 1,000 bootstrap replicates and the GTR+F+ASC+R10 nucleotide model. Additional phylogenetic trees were constructed based on MAFFT v7.310 and Muscle v5.1 (super5 algorithm) with IQ-TREE v2.0.3, FastTree v2.1.10 [79] using the GTR+CAT model, and MEGA v11 [80] using the neighbor-joining method, 1,000 bootstrap replicates and the Maximum Composite Likelihood model. The topologies of resulting trees were manually compared to validate the crucial nodes supporting major conclusions. Phylogenetic trees were visualized using iTOL v6.8 [81].

### Transcriptomic analyses

For transcriptome analyses, a subset of 43 species was selected based on specific criteria: (1) the presence of *FLS*, *DFR*, *F3H*, and *F3’H* genes in the species, (2) the availability of sufficient transcriptomic datasets in the Sequence Read Archive (SRA) and (3) the presence of clear labels for RNA-seq samples with a minimum of three accession numbers corresponding to at least a single plant organ (leaf, root, stem, flower, seed, or fruit). To generate the count tables, all available paired-end RNA-seq datasets of the selected species were retrieved from the SRA (www.ncbi.nlm.nih.gov/sra) [82] using fastq-dump (https://github.com/ncbi/sra-tools). Kallisto v0.44 [83] was then employed to quantify transcript abundance based on the coding sequences of the respective plant species. The previously developed Python script kallisto_pipeline3.py [84,85] served as a wrapper to run kallisto across all input files. Next, the Python script merge_kallisto_output3.py [84,85] was executed to merge the individual kallisto output files into one count table per species. The count tables were processed to create a unified count table, encompassing the expression values of *FLS* and *DFR* across all 43 species. The values of close paralogs were replaced by their sum if these paralogs had the same substrate-preference determining residue at the most important position. RNA-seq based transcript abundance is taken as a proxy for gene expression and thereafter, in the expression analysis, referred to as ‘gene expression’.

To explore the expression patterns of *FLS* and *DFR*, a 2D density heatmap with marginal histograms was generated using the Python script Coexp_plot.py available at https://github.com/bpucker/DFR_vs_FLS.

## Results

### Substrate-preference determining amino acid residues in FLS and DFR and their pattern of occurrence in land plants

From a vast range of diverse taxonomic plants, DFR was identified only in angiosperms. Among the 175 angiosperm datasets analyzed, 129 exhibited at least one most likely functional DFR, characterized by the presence of essential residues associated with DFR activity. Overall, 207 DFR sequences were identified, accounting for cases where multiple DFRs were present in certain species. Multiple sequence alignments of the identified DFR sequences revealed potential NADPH binding and substrate binding sites in N-terminal regions (**Fig 3**). Analysis based on the conservation of the amino acid residue at position 133, i.e., 3rd position within the 26-amino acid-long substrate binding domain of DFR led to the classification of DFR proteins into three types: (i) DFR with asparagine (N) at the 133rd position which can recognize all three dihydroflavonols as substrates, (ii) DFR with aspartic acid (D) at the 133rd position which shows higher preference for dihydroquercetin and dihydromyricetin, and (iii) DFR characterized by alanine (A) at the 133rd position which exhibits a preference for dihydrokaempferol and a higher likelihood of rejecting dihydromyricetin. Henceforth, these three DFR types will be referred to as DFR_N_, DFR_D_, and DFR_A_, respectively.

**Fig 3:**
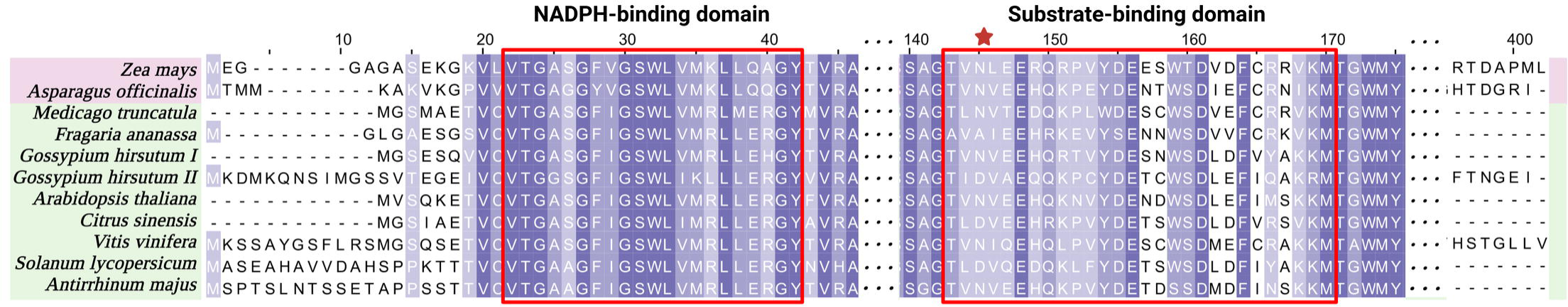
Selected parts of multiple sequence alignment of DFR sequences restricted to 10 species to reduce the complexity. Monocot and dicot species are highlighted by light pink and light green color, respectively. The dark violet, mid violet, and light violet colors indicate 100%, >75%, and >50% similarity among the sequences, respectively. Important domains associated with DFR functionality are highlighted and some columns are masked with three dots (**…**). Red boxes highlight the NADPH-binding domain and the 26 amino acid long substrate-binding domain. The red-star labeled residue at the 3rd position within this region is crucial for dihydroflavonol recognition (position 133 in the *A. thaliana* DFR). MAFFTv7 was applied to generate the alignment.

Sequences of FLS enzymes were successfully identified in the datasets of 143 out of 175 angiosperm species. The inability to identify FLS sequences in some of the plant species may be attributed to technical reasons, i.e., incompleteness of the analyzed data sets. In total, 247 FLS sequences were identified, including instances where multiple copies of FLS were present within a species. FLS enzymes belong to the 2-oxoglutarate-dependent dioxygenase (2-ODD) superfamily. A comprehensive analysis of the multiple sequence alignment revealed the presence of characteristic conserved ferrous iron-binding motif (HX(D/E)X_n_H) and the 2-oxoglutarate binding residues (RXS) critical for FLS functionality (**Fig 4**). Furthermore, sequences of functional FLSs exhibited additional residues (G68, H75, P207, and G261) responsible for the proper folding of the 2-ODD protein. Specific motifs, such as “PxxxIRxxxEQP” and “SxxTxLVP” which were initially considered to be unique to FLS and distinguished it from other plant 2-ODDs [86] were also identified. However, a previous study identified two *BnaFLS1* homologs which also exhibited F3H activity [87]. It was demonstrated that despite having these reportedly FLS-specific motifs, they could still have additional activities. FLS sequences also possessed five key amino acids (H132, F134, K202, F293, and E295) identified as potential DHQ binding site residues, with the latter four exhibiting notable conservation across diverse species. These residues were also found to be conserved in anthocyanidin synthase (ANS) sequences wherein they had a strongly conserved Y132 residue as well. Analysis based on the conservation of residue 132 led to the classification of FLS into three types: FLS_H_ (histidine), FLS_F_ (phenylalanine), and FLS_Y_ (tyrosine). The positions of amino acids are assigned based on the *Arabidopsis thaliana* sequences (*At*DFR, AT5G42800, and *At*FLS1, AT5G08640) for consistency and reference purposes.

**Fig 4:**
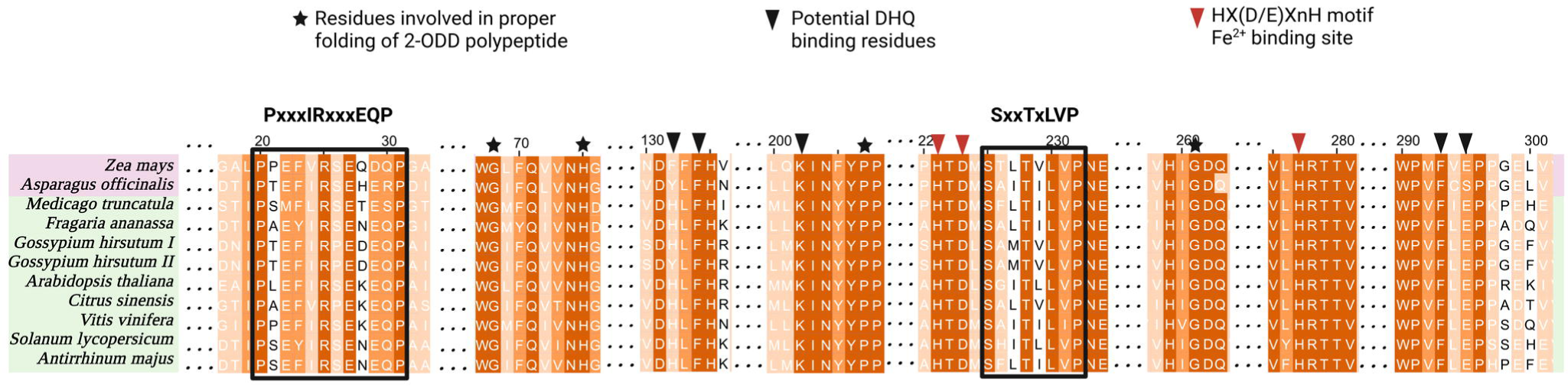
Selected parts of an FLS multiple sequence alignment restricted to 10 species to reduce the complexity. Monocot and dicot species are highlighted by light pink and light green color, respectively. The dark orange, mid orange, and light orange colors indicate 100%, >75%, and >50% similarity among the sequences, respectively. Important domains and residues associated with FLS functionality are highlighted and some columns are omitted as indicated by three dots (**…**). The conserved 2-ODD domain and the Fe^2+^ binding sites are indicated by black asterisks and red arrowheads, respectively. The black boxes highlight the FLS-specific motifs. The black arrowheads indicate potential DHQ-binding sites where the first residue (position 132) is thought to be critical for the substrate preference of FLS. The alignment was generated by MAFFT.

The 129 diverse plant species harboring DFRs represented 28 orders out of the 64 orders recognized in the current Angiosperm Phylogeny Group (APG IV;2016) classification of angiosperms [88]. Interestingly, all monocot plants investigated displayed DFR_N_, with only a few minor exceptions which displayed neither of the three types. Among dicots, some species exclusively possessed either DFR_N_ or DFR_D_, while others exhibited a combination of either DFR_N_ and DFR_D_ or DFR_N_ and DFR_A_. Remarkably, DFR_A_, which is known to have a strong preference for DHK as a substrate, previously reported only in *Fragaria*, was also identified in *Spirodela polyrhiza*, *Spirodela intermedium,* and *Begonia peltatifolia* (Additional File 2). A total of 143 angiosperm plant species encompassing 25 orders were analyzed for functional FLS genes. Among these species, monocots exhibited either FLS_F_ or FLS_Y_, while all dicots possessed FLS_H_ with a histidine residue at position 132. However, certain orders such as Fabales, Fagales, Malpighiales, and Malvales displayed multiple FLS candidates in some species with both FLS_H_ and FLS_Y_, characterized by histidine and tyrosine residues, respectively. (Additional File 3). The overall pattern of amino acid residues at position 133 in DFR sequences and position 132 in FLS sequences at the order level is summarized in **Fig 5**.

**Fig 5:**
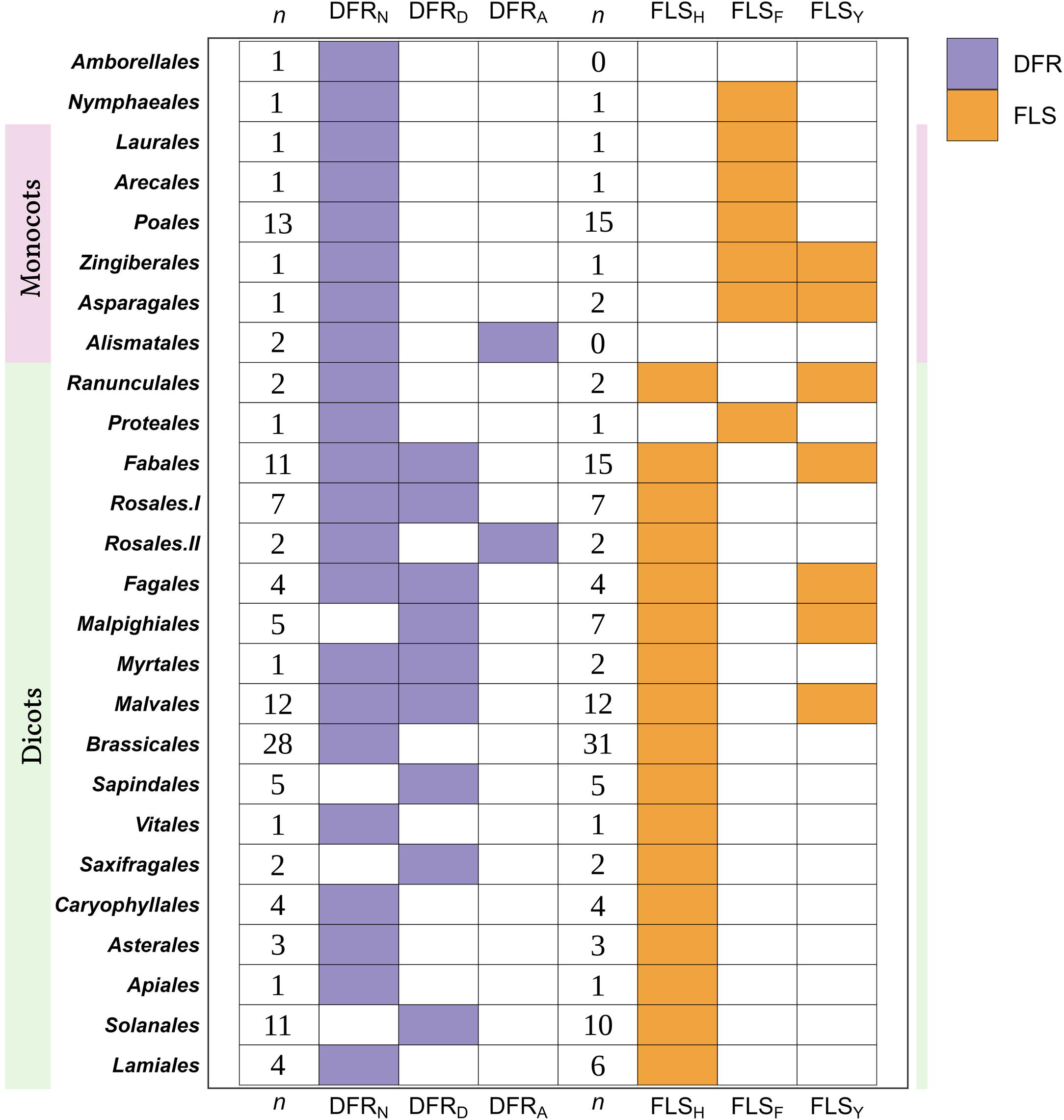
The patterns of commonly occurring amino acid residues at substrate-preference determining positions observed in different orders of angiosperms for FLS and DFR. Orders are sorted by branching in the evolution of angiosperms according to the Angiosperm Phylogeny Group (2016). The residues were investigated across 172 angiosperm species and are represented here at the order level. Monocot and dicot species are highlighted by light pink and light green color, respectively. Rosales II consists of 2 species: *Fragaria x ananassa* and *Fragaria vesca*. *n* represents the number of analyzed species harboring DFR and FLS, respectively, within each order.

### Evolution of DFR

To understand the evolutionary relationships among multiple DFR sequences, a phylogenetic tree based on *DFR* protein-encoding sequences was constructed. Given that DFR belongs to the short-chain dehydrogenase (SDR) family, the closely related ANR, CCR, and DFR-like sequences from this family were included as outgroups. In the resulting analysis, the ANRs, and DFRs clustered together, distinct from the CCRs, as depicted in Additional File 4. The two clades placed as sisters to the CCR clade had *A. thaliana* tetraketide α-pyrone reductase (*AtTKPR1* and *AtTKPR2*) in each clade, essential for pollen wall development and male fertility. TKPR2, known to be more active than TKPR1, clustered closer to the CCR clade. The TKPR1-like, TKPR2-like, CCRs and a large clade with an unknown function clustered together and were placed as a sister clade to the DFR and ANR clade. Adjacent to the DFR clade, another clade featured AtBEN1-encoding a DFR-like protein involved in the brassinosteroid metabolism in *A. thaliana*. This clade did not contain any sequences from the gymnosperms or monocots, only dicots were found. The proximity to the DFR clade and the absence of monocots suggests its evolutionary origin from a DFR predecessor gene after the split of the monocot and dicot lineages.

We did not identify any Gymnosperm DFR in our analysis, possibly due to their early-branching nature and distinct conserved residues compared to angiosperms. Previously characterized DFR sequences (Additional File 5), marked with red asterisks, were generally located within the DFR clade, except for fern *Dryopteris erythrosora*, *DeDFR1* and *DeDFR2*, *Ginkgo biloba*, *GbDFR1*, *GbDFR2*, and *GbDFR3*, and *Camellia sinensis*, *CsDFRc.* The placement of these sequences could suggest an independent origin of DFR activity or enzymatic promiscuity in closely related clades. Fern DFRs, placed outside the ANR and DFR clade, may exhibit distinct characteristics due to their early-branching nature or possess promiscuous functions. Ginkgo DFRs within TKPR-like clades likely deviate from *bona fide* DFRs. The phylogenetic analysis of functional DFRs highlights distinct clades specific to monocots and dicots (**Fig 6**). Monocots mostly exhibit DFR_N_, while dicots generally display various DFR types - DFR_N_, DFR_D_, and DFR_A_. Notably, DFR_D_ appears to have originated from a duplication event of DFR_N_, as observed in the Fabales and Malvales clades. DFR_A_ is restricted in the Rosales and Alismatales clades. These findings suggest possible scenarios where dicots experienced a duplication event of DFR_N_, followed by a subsequent substitution event resulting in the emergence of DFR_D_. Furthermore, in the Rosales and Alismatales clade, the duplication could have been accompanied by a substitution from N to A. Upon examining the codons of the *DFR_A_* coding sequence at position 133 in *F. vesca* and *F. ananassa* and comparison against closely related DFR_N_ sequences, we found that the codons did not exhibit any predisposition towards the emergence of *DFR_A_* in strawberries (AAT to GCC) (Additional file 6).

**Fig 6:**
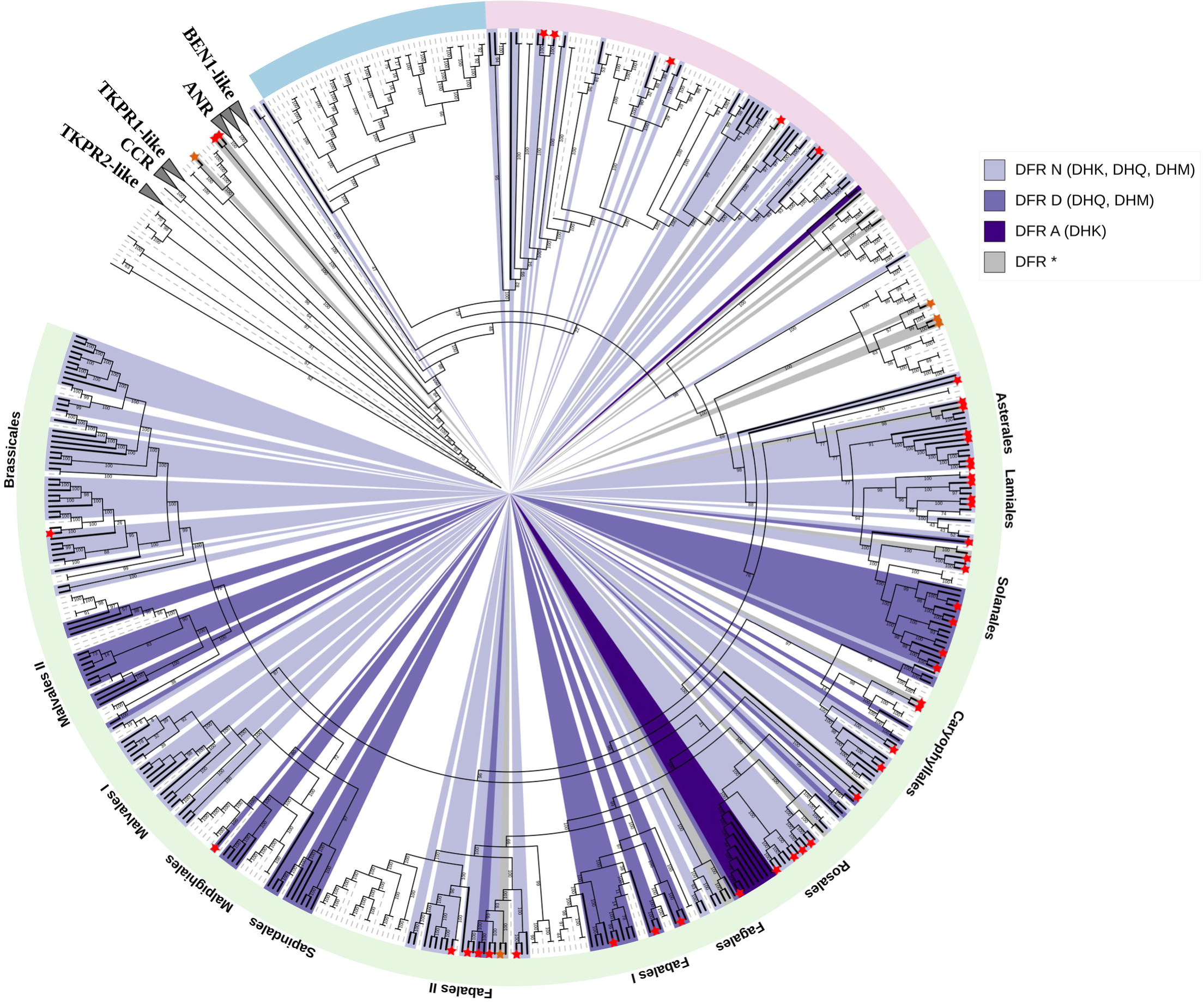
Phylogenetic analysis of DFR sequences in diverse plant species, highlighting amino acid residue diversity at position 133 associated with substrate specificity. Gymnosperm, monocot, and dicot species are denoted by light blue, light pink, and light green color stripes, respectively. Non-DFR sequences are represented by dashed gray branches while the functional DFRs are indicated by solid black branches. The color-coded scheme represents different residues: Asparagine (light purple), Aspartic acid (periwinkle blue), Alanine (deep purple), and other amino acids (gray) at position 133. The preferred substrate of the DFR type is written in brackets: DHK, dihydrokaempferol; DHQ, dihydroquercetin and DHM, dihydromyricetin. Distinct clusters of DFRs from major plant orders are labeled for reference. DFR sequences identified in previous studies are highlighted by an asterisk at the start of the terminal branch, with asterisks of functional DFR genes colored in red (Additional File 5). Leaf labels are hidden to reduce the complexity. The outgroup comprises SDR members like ANRs, CCRs, and other DFR-like sequences (Additional File 4).

### Deep duplication events during the evolution of FLS

Through a phylogenetic tree generated with *FLS* protein-encoding sequences, we observed deep duplication events during the evolution of FLS enzymes. Considering their classification within the 2-oxoglutarate-dependent dioxygenase (2-ODD) family, closely related ANS, F3H, and FNSI sequences from the same family and FLS-like sequences lacking crucial residues for FLS functioning were included as outgroups. The phylogenetic tree revealed a largely distinct clustering of F3H, ANS, and FLS sequences. ANS and FLS formed a common cluster, while F3H grouped with FNSI sequences (Additional File 7).

The non-Apiaceae FNSI sequences clustered outside the F3H clade while the liverworts FNSI and Apiaceae FNSI were a part of a single clade with phylogenetically distinct F3H sequences. The liverworts FNSI are present at the root of this clade suggesting that plant F3H evolved from liverworts FNSI. An independent lineage of Apiaceae FNSI is located nested within the Apiaceae F3H sequences further confirming the evolution of Apiaceae *FNSI* from *F3H* [84,89]. Gymnosperm, monocot, and dicot sequences formed distinct clades within larger clades corresponding to the three 2-ODD members, i.e., F3H, ANS, and FLS. The FLS clade was divided into four primary clusters: a Gymnosperm clade branching first, a second clade which we hereafter call aFLS (ancestral FLS), a third monocot clade, and lastly, a large dicot clade. The aFLS clade encompasses FLS-like sequences from basal angiosperms and dicots and a few dicot FLS sequences. Only nine FLS sequences, one each from *Linum usitatissimum, Lotus japonicus, Cajanus cajan, Quercus suber, Betula platyphylla, Acacia confusa AcFLS* (JN812062), *Camellia sinensis CsFLSa* (KY615705), *Fagopyrum tataricum FtFLS2* (JX401285), *and Vitis vinifera FLS5* (AB213566) were in this clade. Although these FLS-like sequences covered a wide range of angiosperms, no monocots were represented in this clade. The presence of this aFLS clade sister to the distinct monocot and dicot clade implies an FLS duplication event preceding the monocot-dicot split. The large dicot cluster further separates into two distinct subclades: one including sequences exclusively belonging to the Brassicales order, and the other including sequences from the rest of the dicot plants (**Fig 7**). In the Brassicales subclade, a monophyletic group of Brassicales FLS and FLS-like sequences is evident, with *AtFLS1* and sequences of other functional Brassicales FLSs clustered together. The presumed non-functional FLS sequences from the Brassiclaes order group together and are classified based on their phylogenetic relationship to their most likely *A. thaliana* orthologs (*AtFLS2*-*AtFLS6*). Within the latter subclade which excludes Brassicales, some FLS sequences from the same order appear in distinct clusters as observed in the Fagales, Sapindales, Malpighiales, and Malvales order, suggesting gene duplication events in the evolutionary history of FLS in some common ancestor of these orders.

**Fig 7:**
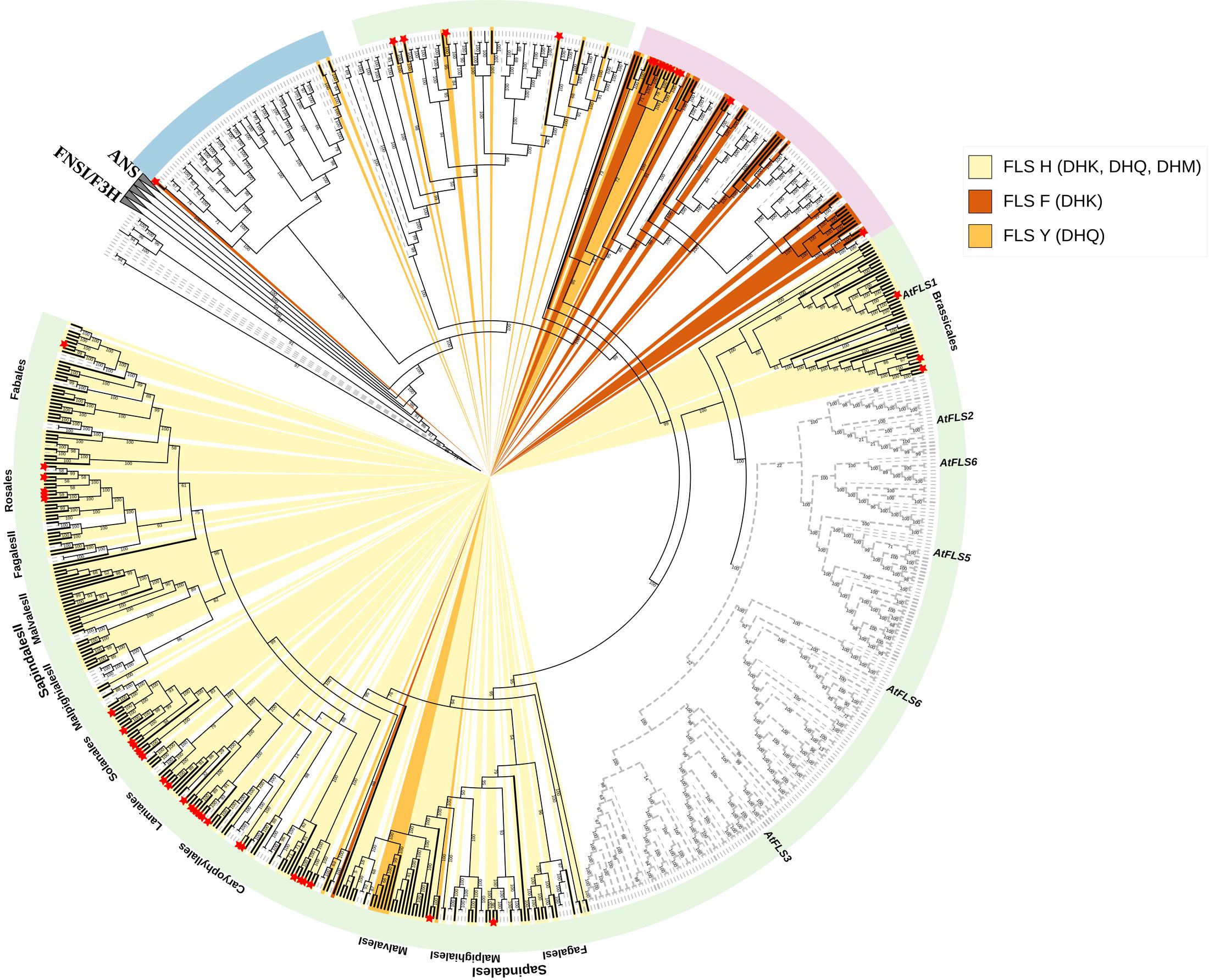
Phylogenetic diversity of FLS sequences in diverse plant species, highlighting amino acid residue diversity at position 132 associated with DHQ-substrate binding. Gymnosperm, monocot, and dicot species are denoted by light blue, light pink, and light green color stripes, respectively. Non-FLS sequences are represented by dashed gray branches while the functional FLSs are indicated by solid black branches. The background color highlights different presumably substrate-preference-determining amino acid residues: histidine (pale yellow), phenylalanine (dark orange), and tyrosine (dark golden) at position 132. The hypothesized preferred substrate of the FLS type is written in brackets: DHK, dihydrokaempferol; DHQ, dihydroquercetin, and DHM, dihydromyricetin. Distinct clusters of FLSs from major plant orders are labeled for reference. FLS sequences identified in previous studies are highlighted by an asterisk at the start of the terminal branch with asterisks of functional FLS genes colored in red (Additional File 5). The branches of *Arabidopsis thaliana* AtFLS1-AtFLS6 are labeled. Individual leaf labels are hidden to reduce the complexity. The outgroup comprises members of 2-ODD like F3H, ANS, and other FLS-like sequences (Additional File 7).

While the first and last residues of the FLS substrate binding residues (H132, F134, K202, F293, and E295) exhibit diverse residues, the central three are strongly conserved in both FLS and ANS sequences. Notably, gymnosperm FLSs and aFLSs feature a Y132 residue, consistent with ANS sequences. Monocotyledonous plants display an F/Y(85%/15%) residue at position 132, whereas the majority of dicots consistently possess an H residue at position 132, except for multiple *Gossypium* species and *Herrania umbratica* from the Malvales order, which have a tyrosine residue at this position. The E295 residue is conserved in ANS sequences as well as in monocot and dicot FLSs. However, sequences from the gymnosperm, aFLSs, AtFLS4, and AtFLS5 clades exhibit non-conserved amino acid residues at position 295. The aFLSs exclusively feature aliphatic amino acids at this position (G/A/V/L/I).

### Divergent expression patterns of *FLS* and *DFR*

Generally, either *DFR* or *FLS* is expressed in a given sample, but almost never both simultaneously (**Fig 8a**). This observation held true across various *FLS* and *DFR* types, underscoring a general phenomenon. However, the pattern intriguingly deviates in some cases. The almost mutually exclusive expression patterns were apparent for all combinations of *FLS* and *DFR* types (**Fig 8b-g**). These patterns were particularly pronounced in the *DFR_A_* vs *FLS_H_* expression and *DFR_N_* vs *FLS_Y_* expression plots. Contrastingly, in cases such as *DFR_N_* vs *FLS_H_* and *DFR_D_* vs *FLS_H_*, instances of co-expression of *FLS* and *DFR* were observed in some samples. Theoretically, there could be 9 different *FLS-DFR* combinations, given three *FLS* and three *DFR* types were examined. The absence of certain combinations raises the possibility that not all *FLS* and *DFR* combinations may occur in nature. For instance, to the best of our knowledge, *FLS_F_* is restricted to monocots, exclusively paired with *DFR_N_*. Hence, combinations involving *DFR_D_* and *DFR_A_*alongside *FLS_F_* might not exist in nature.

**Fig 8:**
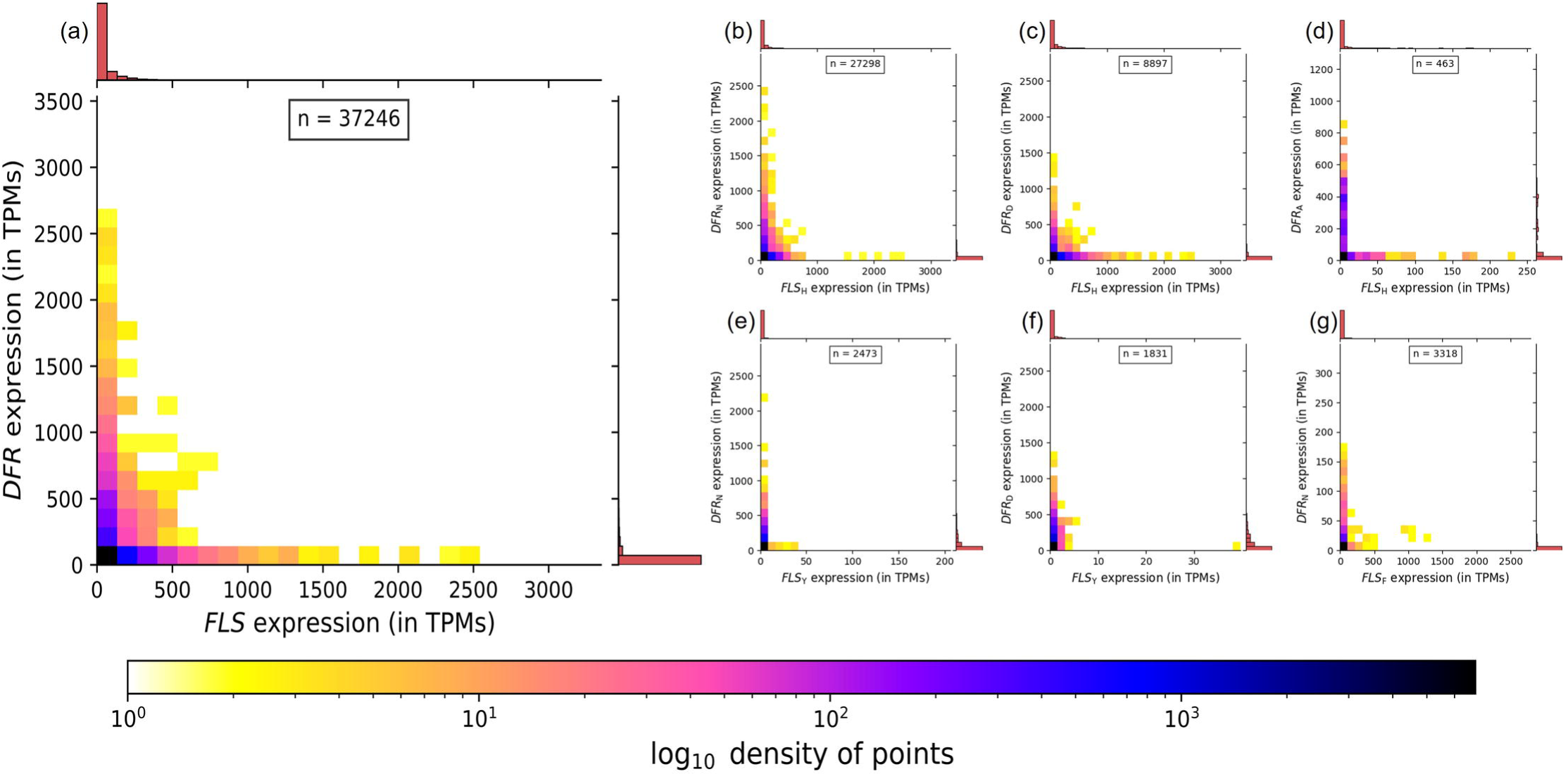
2D density heatmap with marginal histograms showing divergent expression patterns of FLS vs DFR across samples from 43 species. Each sample shows only the expression of *FLS* or *DFR*, but not both. The sample size is indicated by n. Colorbar depicts the logarithmic density of points in the plot. (a) Combined expression of all *FLS* and *DFR* types, (b-g) specific combinations of *FLS* and *DFR* types.

Besides *FLS* and *DFR*, the major genes influencing the dihydroflavonol levels are *F3H*, *F3’H*, and *F3’5’H*. These genes play a crucial role in determining the hydroxylation patterns of dihydroflavonols, which are common substrates for FLS and DFR (**Fig 1**). We analyzed the tissue-specific expression of *F3H*, *F3’H*, and *F3’5’H* in 43 species and correlated the expression data with different DFR and FLS types (Additional File 8, Additional File 9). The results suggest that the production of specific dihydroflavonols is more active in some species compared to others. The heatmap analysis reveals distinct expression patterns: plants like *Cicer arietinum* and *Citrus sinensis* exhibit high FLS activity, while *Vitis vinifera* and *Dianthus caryophyllus* show higher DFR activity. The red leaf of *Lactuca sativa* displayed higher expression of *F3H*, *F3’H*, and *DFR_N_* compared to the green leaf, indicating a greater substrate availability in the red leaf. Differences in gene expression are linked to substrate preferences and pigment production, impacting plant coloration (Additional File 8).

## Discussion

Factors influencing the competition of the pivotal enzymes FLS and DFR for the common substrate, dihydroflavonols, remain poorly understood. The equilibrium between these enzymes can be modulated by several mechanisms including (i) the unique substrate specificity exhibited by FLS and DFR for different dihydroflavonols, (ii) the existence of multiple FLS/DFR copies with narrow substrate preferences, or (iii) the expression of specific *FLS*/*DFR* copies in specific cell types or under specific conditions, (iv) the availability of substrates, i.e., different dihydroflavonols, determined by the activity of flavonoid 3’-hydroxylase and flavonoid 3’,5’-hydroxylase, and (v) the channeling of the substrate through metabolons.

### Mitigating substrate competition between FLS and DFR

A well-studied 26-amino acid region determines substrate specificity in DFR [55,90] and the grape crystal structure further emphasized the significance of the N133 amino acid in binding the 3’ and 4’ hydroxyl groups of DHQ [63]. Compared to DFR, the substrate specificity determining residues of FLS has received less attention. The substrate-binding residues of FLS (H132, F134, K202, F293, E295) show high conservation among all plant FLSs, except the H132 and E295 residues. Through the examination of *A. thaliana fls1* mutant, it was hypothesized that the H132 residue may play a critical role in governing the diverse substrate preferences of FLS enzymes [64]. Numerous previous studies have employed enzyme assays to investigate the substrate preferences of plant FLSs and DFRs, while many others identified the type of anthocyanins and flavonols in various plants. Metabolite accumulation within a system could be influenced by factors beyond substrate specificity like enzyme abundance, cellular compartmentalization, and metabolic regulation. Hence, to evaluate the general substrate specificity of an enzyme, it is best to compare *k*_cat/_*K*_M_ values rather than metabolite accumulation. Due to the unavailability of enzyme kinetics data in previous studies, we had to utilize metabolite accumulation as a proxy to derive substrate preferences of DFR and FLS. For instance, *AcFLS* from the monocot *Allium cepa*, characterized by the Y132 residue, exhibits a preference for DHQ over DHK [91], while monocots such as *OsFLS* [92] and *ZmFLS* [93], with the F132 residue, favor DHK over DHQ. In contrast, DFR_N_ in maize and *Dendrobium officinale* predominantly produces cyanidin derivatives [94,95]. DFR_N_ of barley displays *in vitro* reduction of DHK and DHQ whereas DHM was not tested [96]. Reports of procyanidin and prodelphinidin in testa tissue and a lack of detection of leucopelargonidin and its derivatives [97] do not align with a DHK and DHQ specificity of DFR in barley. DFR_N_ of grape hyacinth and *Cymbidium hybrida* exhibit high activity towards DHM and DHQ [90,98], and *Anthurium andraeanum* DFR, featuring an uncommon S133 residue, displays a marked specificity for DHK [99].

Previous studies and our analyses identified multiple DFR copies in dicots. For example, in the Fabales order, *M. truncatula* DFR1 and *L. japonicu*s DFR2 and DFR3 with an N133 residue reduces DHK more readily than DHQ, while *M. truncatula* DFR2 and *L. japonicus* DFR5 with a D133 residue show a higher activity towards DHQ [61,62]. In these studies, the relative DFR activity was demonstrated by HPLC and spectrophotometric assays, respectively. In the Rosales order, FLS_H_ from *Fragaria ananassa* [100], *Malus domestica* [101], and *Rosa hybrida* [102] reduces both DHK and DHQ, with strawberry FLS favoring DHK and rose FLS favoring DHQ. Conversely, their DFR_N_ exhibits a higher affinity for DHQ than DHK [68,103,104]. The strawberry species, *Fragaria ananassa* and *Fragaria vesca* have another DFR-type, DFR_A_ which has a very high specificity for DHK [68]. Other examples are *L. japonicus* and *M. truncatula*, where a DFR_D_ is present with a high preference for DHQ, while the DFR_N_ prefers DHK.

This observation seems to contradict previous reports about the substrate preferences of DFR_N_. A potential explanation might be sub-/neofunctionalization upon duplication of DFR. We speculate that through a yet unknown mechanism, DFR_N_ might become more specific for the substrate that is not favored by the additional DFR copy. Waki et al. reported that chalcone isomerase-like proteins (CHILs) can bind to chalcone synthase (CHS) and rectify its promiscuous activity making CHS more specific [105]. In a conceptually similar mechanism, multiple DFR copies with distinct specificities could be influenced by external factors to adjust their substrate preferences, i.e., to make the DFR_N_ type accept the substrate not favored by the additional DFR copy.

In the Brassicales, FLS_H_ from *A. thaliana* and *Matthiola incan*a prefer DHK [106–108], while their DFR_N_ reduces both DHK and DHQ but favors DHQ [109–111]. *Citrus unshiu* FLS_H_ has higher DHK activity [112], whereas *Zanthoxylum bungeanum* from the same order, Sapindales, has DFR_D_ and prefers DHM followed by DHQ and DHK [113]. In Caryophyllales, FLS_H_ in *Dianthus caryophyllus*, *Fagopyrum dibotrys*, and *Fagopyrum tataricum* catalyze both DHK and DHQ [114–116], while DFR_N_ in *Dianthus* reduces DHQ more effectively than DHK [115]. In *Fagopyrum esculentum*, DFR_N_, active in all tissues, reduces both DHK and DHM at the same rate, while the second DFR enzyme with a V134 residue, expressed only in roots and seeds, exhibits high specificity towards DHQ [67]. In Solanales, *Petunia hybrida* and *Nicotiana tabacum* FLS_H_ catalyze both DHK and DHQ with minimal activity towards DHM [58,117], whereas their DFR_D_ is specific in reducing 3,4 di- and 3,4,5 tri-hydroxylated substrates but apparently cannot reduce DHK [55,56]. *Gentiana triflora* predominantly produces blue flowers and generally has cyanidin, delphinidin, and quercetin derivatives [118].

In our study, we observed that all monocots were mostly FLS_F_ type, and some were FLS_Y_ type, while their DFR were predominantly DFR_N_ type. Enzyme assay studies suggest that in monocots, FLS_F_ generally prefers DHK while DFR_N_ prefers double or triple-hydroxylated dihydroflavonol substrates. This difference in substrate preferences could help in avoiding competition between the two enzymes. In dicots, FLS_H_ is observed in the majority of plants, but in fewer plants with multiple FLS sequences, FLS_Y_ is also observed. While DFR is more diverse in dicots, some have only DFR_N_ or only DFR_D_, while others have DFR_N_ along with either DFR_D_ or DFR_A_ candidates. Prior studies collectively suggest that FLS_H_ in dicots can utilize both DHK and DHQ as substrates but may have a preference for one over the other in different plants, implying that the H132 residue remains more neutral between DHK and DHQ. The biosynthesis of DHM requires the presence of flavonoid-3’,5’-hydroxylase (F3’5’H) activity, which is rare. Dyer et al suggested that the competition for pollinators might have driven the evolution of blue flower color [119,120]. DHQ and DHM (if *F3’5’H* is present) appear to be the preferred substrate of DFR in most cases. Both in monocots and dicots, double and triple-hydroxylated substrates seem to be preferred by DFR, aligning with the notion that cyanidin is the most abundant and widely distributed floral anthocyanin [121]. While substrate-specifying residues may play a role in identifying the specificity of FLS and DFR enzymes, it is plausible that substrate preference may not be exclusively determined by the H132 and N133 residues, respectively, in all plants. Given the rapid progress in protein structure prediction [122], it might be possible to quantify the contribution of additional residues in the near future through the integration of all available data.

### Functional divergence during DFR evolution

Short-chain dehydrogenases/reductases (SDR) is an NADP-dependent enzyme superfamily ubiquitous across all life forms. DFR, belonging to the SDR108E enzyme family, converts dihydroflavonols (DHK, DHQ, and DHM) into colorless leucoanthocyanidins. These leucoanthocyanidins subsequently undergo further modifications to produce a wide array of pigments such as anthocyanins and proanthocyanidins (also known as tannins). Cinnamoyl-CoA reductases (CCRs), involved in lignin biosynthesis, likely originated early in land plants and diverged distinctly from other SDR108E family [137], as seen by the distinct clustering of CCRs with DFR and ANR in our phylogenetic analysis. *DeDFR1* and *DeDFR2* in *Dryopteris erythrosora*, *GbDFR1*, *GbDFR2*, and *GbDFR3* in *Ginkgo biloba*, and *CsDFRc* in *Camellia sinensis* were positioned outside the clade comprising functionally characterized DFRs. Fern DFRs, showing both *in vitro* and *in vivo* DFR activity, displayed a unique R residue at position 133, exhibiting substrate specificity for DHK and DHQ but an inability to catalyze DHM [138]. The positioning of these fern DFRs outside the DFR and ANR clade could suggest the independent evolution of DFRs in ancient land plants. Notably, ginkgo DFRs [139], questioned by Halbwirth et al (2019) [48] for their authenticity, were found within TKPR-like clades in our analysis, suggesting their classification as DFR-like sequences rather than *bona fide* DFRs. *CsDFRc* capable of restoring *Arabidopsis tt3* mutant phenotypes [129] supports its DFR activity. At first glance, the positioning of CsDFRc outside the functional DFR clade could suggest that it might have evolved independently. Further investigation revealed that the clade containing *CsDFRc* also harbored an *Arabidopsis* sequence, AT4G27250, named ABA HYPERSENSITIVE 2 (ABH2) in a study by Weng et al (2016) [140]. This ABH2 is the long-sought-after phaseic acid reductase (PAR), which is a DFR-like enzyme involved in abscisic acid catabolism. This suggests potential promiscuous functions of the sequences in this clade where they have a PAR functionality but could also catalyze DFR-like reactions, warranting further exploration in future studies. A thorough examination of the clade comprising functionally characterized DFRs revealed primarily three distinct DFR types determined by the amino acid residue at position 133 - DFR_N_, DFR_D_, and DFR_A_. Our phylogenetic analysis indicates that DFR underwent duplication events, likely associated with N133D and N133A substitutions. These duplications and substitutions led to the acquisition of novel or enhanced functions, characterized by narrower substrate specificity, compared to the non-specific DFR_N_. Until now, DFR_A_ was considered exclusive to the *Fragaria* genus, but our detailed exploration has revealed DFR_A_ in the *Spirodela* genus (*S. intermedium* and *S. polyrhiza*) and *Begonia peltafolia*. This discovery suggests a more widespread distribution of DFR_A_ and multiple independent origins, although the power of this study is limited by available plant sequences in public repositories.

While we could not identify DFR in gymnosperms, the DFR sequences of monocots and dicots formed distinct clades. Notably, two monocot clades were observed—one dominated by DFR_N_ sequences and another smaller one situated at the root of the dicot clade, harboring non-DFR_N_ sequences of monocots. These sequences exhibited amino acid diversity at position 133, such as cysteine (C) in *Musa acuminata*, alanine (A) in *Spirodela polyrrhiza*, and t*hreonine* (T) in *Dioscorea alata* and *D. rotundata*. While cysteine and threonine share similar properties with alanine, predictions about their performance, such as a preference for DHK, remain speculative. Given the lack of enzyme assay studies for these proteins, the functional status of this clade remains an open question.

Our analysis identified DFRs that do not conform to DFR_N_, DFR_D_, or DFR_A_ types, albeit in limited numbers. Few such DFRs have already been described, like *Lotus japonicus DFR1* [62], and *Ipomoea nil DFR-A* [141], featuring residues S and H at position 133, respectively, were found to lack DFR activity. It remains unknown whether these have undergone neo-functionalization or are pseudogenes. Others, like monocot *Anthurium andraeanum* DFR, with a S133 residue, exhibit high activity towards DHK [99], while *Vaccinium macrocarpon DFR1-1*, *DFR2-1* [142], and *Fagopyrum esculentum FeDFR2* [67], featuring a V133 residue, prefer DHQ over other dihydroflavonols. Additionally, we identified DFR sequences with residues not studied by Johnson et al, including *Musa acuminata* (C133), *Malus domestica* (I133), *Carica papaya* (S133), *Fagus sylvatica*, and *Quercus suber* (R133). Investigating residues beyond N, D, and A at position 133 could unveil further insights into DFR functionality and substrate specificity.

DFRs with known functional activity and specificity towards DHK, as observed in strawberries where DFR_A_ is active in a ‘false fruit,’ and in A. andraeanum where DFR_S_ is active in spathe leaf— a modified leaf—suggest that unconventional or modified structures could prompt the evolution of different DFRs. Additional DFR types may be discovered in the future within specific organs of plants that assume the function of fruit, flower, or leaves.

### Functional divergence during FLS evolution

In the plant kingdom, the 2-oxoglutarate-dependent dioxygenase (2-ODD) superfamily is one of the largest enzyme families only second to cytochrome P450s (CYPs). Within this superfamily, the plant 2-ODD family can be categorized into three classes: DOXA, DOXB, and DOXC, with all enzymes involved in flavonoid biosynthesis falling into DOXC [123]. These 2-ODD enzymes are believed to have originated from a common ancestor before the emergence of land plants, and then underwent species-specific evolution in response to diverse environmental conditions [123]. The evolution of FLS and ANS is likely to have occurred after the emergence of F3H during seed plant evolution [123]. A noteworthy observation similar to that reported by Wang et al. (2019) [124], is the placement of ginkgo 2-ODD genes (*GbFLS*, *GbANS*, and *GnF3H*) into FLS, ANS, and F3H clades, respectively, indicating the divergence of F3H, FLS, and ANS preceded the separation of gymnosperms and angiosperms.

A comprehensive analysis of the functional FLS clade revealed a deep duplication in the evolutionary history of the FLS gene. We found that the FLS could be split into four different subgroups, gymnosperm sequences, aFLS, monocot sequences, and dicot sequences. The aFLS sequences consisting of dicot sequences were more similar to gymnosperm FLSs than to the monocot and dicot FLS sequences in the distinct latter clades suggesting the duplication of ancestral FLS before the divergence of monocots and dicots. The aFLS clade contains FLS-like sequences from basal angiosperms and dicots and some previously characterized dicot FLS sequences. They have a Y132 residue and an aliphatic amino acid residue at position 295, mostly valine. Some of the FLS sequences in the aFLS clade are previously described. For example, the *Fagopyrum tataricum FtFLS2* is mainly expressed in roots, flowers, and immature seeds and is upregulated by exogenous application of salicylic acid and NaCl and not affected by abscisic acid [125–127]. However, activity analysis of *FtFLS2* was not performed. *Camellia sinensis CsFLSa* has high expression in buds but the buds mostly accumulate catechins. Heterologous expression of *CsFLSa* in tobacco failed, hence, FLS functionality remains unknown [128]. In *Vitis vinifera, FLS5* has very high transcript levels but the quercetin accumulation coincides with the transcription of *FLS4* (H132 residue) and not *FLS5*. The shaded berries accumulate as much *FLS5* mRNA as the control berries while the *FLS4* (H132 residue) mRNA is not accumulated at all [129]. *AcFLS* is expressed in almost all plant parts and the highest abundance is seen in flowers. The maximum level of *AcFLS* mRNA level was recorded six hours after wounding [130]. We observed in our study that *Lotus japonicus FLS4* has a low overall transcript level. No previously described aFLS sequence has been tested to show *in vivo* FLS activity. The position of aFLS clade separate from the canonical FLS clade could indicate that members of the ancestral FLS clade might not be canonical FLS, but have undergone potential neofunctionalization since separation from the canonical FLS lineage.

The large dicot clade comprises a Brassicales order subclade and another clade comprising the rest of the dicots. Both clades show multiple duplication patterns at a shallower level. We presume that sequences in the Brassicales clade have functions similar to their *A. thaliana* AtFLS1-AtFLS6 orthologs. Preuss et al (2009) demonstrated that only *At*FLS1 and *At*FLS3 exhibit FLS activity, with other AtFLSs identified as non-functional and AtFLS3 only showing activity under extended assay conditions [131]. However, other studies only found strong evidence for FLS activity in the Brassicales FLS1 clade [87,106]. The presence of the FLS2-FLS6 clade across various Brassicales species suggests a potential undiscovered function for these enzymes. Distinct E295 residue patterns are observed among the FLS clades; AtFLS1, AtFLS2, AtFLS3, and AtFLS6 have E/D295 residue, whereas AtFLS4 and AtFLS5 have other non-conserved residues at this position. The AtFLS5 clade features aliphatic amino acids at this position, and *AtFLS5* shows expression primarily in the roots - especially in the roots of seedlings [106]. Enzyme assay results by Chua et al. (2018) indicate that the mutations H132Y and H132F show higher specific activity than wild-type *Arabidopsis* FLS1, with E295L mutation resulting in only 7% of WT activity. However, the study only used DHQ as a substrate, and DHK was not tested. Additional studies testing the enzyme activity of mutant FLS (H132Y, H132F, and E295(V/L)) with different substrates would help to confirm the function of these aFLSs and the substrate preferences of FLSs with different residues.

In the second subclade, an interesting evolutionary pattern is observed, where FLS sequences from some plant orders appear in two different clades, particularly noticeable in the Fagales, Malpighiales, Malvales, and Sapindales orders. This suggests a gene duplication during the evolution of land plants probably before the split of the aforementioned orders followed by a significant divergence in FLS sequences in these orders due to functional requirements, leading to the formation of distinct FLS clades. Within the Malvales order, including cotton species, an additional duplication event was observed, originating from FLS_H_ duplication, followed by a subsequent substitution leading to the re-emergence of FLS_Y_ (H132Y). The exclusive presence of the Y residue at position 132 in the Malvales order indicates a potential evolutionary step to enhance substrate affinity. Other orders like Solanales, Rosales, and Fagales are only represented in a single FLS clade.

Although the *Citrus unshiu* FLS within the Sapindales I clade exhibits a significantly greater affinity for DHK compared to DHQ, it predominantly accumulates quercetin 3-O-rutinoside (rutin) [132]. The authors of this study proposed the presence of more than one FLS in *C. unshiu* [112]. In other *Citrus* species, like *C. sinensis*, *C. trifoliata,* and *C. clementin*a, we observed two FLS candidates, one each in the Sapindales I and II clade. Our analysis and enzyme assay data from previous studies suggest that FLSs within the first dicot subclade (Fagales I, Sapindales I, Malpighiales I, and Malvales I) display a pronounced substrate specificity for DHK. In contrast, FLS sequences within the second dicot subclade show a preference for both DHK and DHQ and may have a higher specificity for DHQ [100,102,114,115,128,129,133].

ANS/LDOX, which catalyzes the step immediately downstream of DFR, is closely related to FLS and shows 50-60% polypeptide sequence similarity. Besides the conserved 2-ODD family residues, ANS/LDOX and FLS have similar amino acid residues that have been implicated in the DHQ binding of FLS (Y132, F134, K202, F293, and E295 conserved ANS/LDOX residues based on position in AtFLS1) and show the same substrate/ascorbate binding residue, E230 in AtLDOX. Bifunctional ANS enzymes with FLS-like activity have been discovered in *G. biloba* [134], *O. sativa* [135], *A. thaliana* [65], and *M. truncatula* [136]. *In vitro* studies have shown that ANS can convert leucocyanidin to cyanidin and also dihydroquercetin to quercetin. *At*LDOX [131] converted DHQ and DHK into quercetin and kaempferol, respectively, in the ratio of 1:0.7 suggesting a higher affinity for DHQ at least in *A. thaliana*. In the case of flavonols, FLS might be specific for DHK and ANS might be responsible for converting DHQ and DHK (to a lesser extent) to respective flavonols. This could be the reason behind different patterns of kaempferol and quercetin accumulation observed in different plants. In the case of anthocyanidins, DFR is generally more likely to accept DHQ and the ANS also might be more adapted to produce leucocyanidins.

### *FLS* and *DFR* expression pattern contributes to competition mitigation across angiosperms

The exploration of gene expression across 43 angiosperm species revealed global patterns and offered insights into the competition mitigation between FLS and DFR. There is a predominant concentration of data points along the x and y axes (Fig 8), indicating that co-expression of FLS and DFR almost never occurred. This observation suggests that in the competition between FLS and DFR, only one of them is active, directing the substrate either towards flavonol or anthocyanins. The separation might even take place at the single-cell level, which could explain samples that show some activity of both genes. The analyzed RNA-seq samples are mixtures of different cell types thus FLS and DFR expression could be divergent, but would appear as co-expression because RNA from multiple different cell types is mixed during extraction. Transcription factors might be responsible for causing an almost mutual exclusion of FLS and DFR expression. For example, the presence of DFR_A_, known for preferring DHK, alongside FLS_H_, also inclined towards DHK, was associated with the exclusive expression of only one of the corresponding genes within each sample. Similar dynamics were observed for DFR_N_ and FLS_Y_, where both might favor DHQ, providing empirical evidence for substrate competition dynamics. In other cases, like FLS_H_ and DFR_N_ or FLS_H_ and DFR_D_, there are some samples where both genes are co-expressed. The preference of the above-mentioned DFRs for DHQ and DHM, juxtaposed with FLS_H_’s affinity for DHK emphasizes the intricate substrate-specific regulation between the genes, thereby mitigating the competition.

### Power and limitations of big data upcycling

The wealth of available transcriptomic data offers unprecedented opportunities for efficient data upcycling and comprehensive cross-species comparison. However, it is crucial to acknowledge that many datasets in the Sequence Read Archive (SRA) lack comprehensive metadata, and in some instances, samples may suffer from cryptic or mislabeling issues, posing challenges for data upcycling [143]. Therefore, careful consideration, filtering, and validation of sample tissue information are imperative to ensure the reliability and robustness of data interpretation and subsequent analyses. Nonetheless, the benefits of data reuse in biology outweigh the challenges, especially with the surge in publicly available data and the affordability of sequencing [144]. In this study, we harnessed transcriptomic and genomic data from 43 species distributed across the angiosperms to perform expression analyses. We also utilized vast amounts of genomic data available for a large number of species and employed tools like KIPEs3 to identify functional flavonoid pathway genes with high specificity and accuracy [73]. Achieving this through wet lab methods would not only be laborious, time-consuming, and costly, but also impossible due to the unavailability of the biological material. This underscores the importance of open data to enable efficient exploration of complex biological processes and to facilitate the discovery of novel insights.

## Conclusion

The branching point of the flavonoid biosynthesis into flavonols, catalyzed by FLS, and anthocyanins/proanthocyanidins, catalyzed by DFR, is a pivotal step for metabolic flux control. The availability of massive plant sequence datasets and gene expression resources enabled the investigation of FLS and DFR across a wide spectrum of plant species. This transformative approach revealed global patterns that cannot be identified by analyzing individual gene functions in one species through classical wet lab methods. Amino acid residues associated with substrate preferences, evolutionary patterns, and potential competition mitigation mechanisms were explored. These insights into competing branches responsible for producing colored anthocyanins and non-colored flavonols can guide molecular breeding or engineering of ornamental plants.

## Supporting information

AdditionalFile1

AdditionalFile2

AdditionalFile3

AdditionalFile4

AdditionalFile5

AdditionalFile6

AdditionalFile7

AdditionalFile8

AdditionalFile9

## Declarations

### Data Availability

Scripts developed for this project and additional datasets are available on GitHub (https://github.com/bpucker/DFR_vs_FLS). The count tables of all 43 species analyzed via RNA-seq are freely available through LeoPARD (https://doi.org/10.24355/dbbs.084-202306231402-0) [145].

### Authors’ contribution

NC and BP designed the experiments, collected data, wrote scripts for the data analysis, generated figures, and wrote the manuscript. All authors read and approved the final version of this manuscript.

## Acknowledgments

We thank the research group Plant Biotechnology and Bioinformatics at TU Braunschweig for the excellent discussions. This work was supported by the BMBF-funded de.NBI Cloud within the German Network for Bioinformatics Infrastructure (031A532B, 031A533A, 031A533B, 031A534A, 031A535A, 031A537A, 031A537B, 031A537C, 031A537D, 031A538A). We would like to thank the entire de.NBI team for their excellent technical support. bioRender.com was used to construct some of the figures. We acknowledge support by the Open Access Publication Funds of Technische Universität Braunschweig.

## Additional Files

**Additional File 1:** References of the coding sequence data sets that were used in this study.

**Additional File 2:** Collection of ANR, CCR, DFR, and DFR-like sequences that were used for the phylogenetic analysis of DFR.

**Additional File 3:** Collection of F3H, FNSI, ANS, FLS, and FLS-like sequences that were used for the phylogenetic analyses of FLS.

**Additional File 4:** Phylogenetic trees of DFR. (1) DFR tree showing outgroups constructed by IQ-TREE based on an MAFFT alignment, (2a-f) DFR-specific trees with collapsed outgroups. Gymnosperm, monocot, and dicot species are denoted by light blue, light pink, and light green color stripes, respectively. Non-DFR sequences are represented by dashed gray branches while the functional DFRs are indicated by solid black branches. The color-coded scheme represents different substrate-preference-determining residues at position 133: Asparagine (light purple), Aspartic acid (periwinkle blue), Alanine (deep purple), and other amino acids (gray). The preferred substrate of the DFR type is written in brackets, DHK, dihydrokaempferol; DHQ, dihydroquercetin and DHM, dihydromyricetin. Distinct clusters of DFRs from major plant orders are labeled for reference. DFR sequences identified in previous studies are highlighted by an asterisk at the start of the terminal branch, with asterisks of functional DFR genes colored in red. (2a) Constructed by IQ-TREE based on a MAFFT alignment, (2b) constructed by IQ-TREE based on a Muscle5 alignment, (2c) constructed by FastTree2 based on a MAFFT alignment, (2d) constructed by FastTree2 based on a Muscle5 alignment, (2e) constructed by MEGA based on a MAFFT alignment, and (2f) constructed by MEGA based on a Muscle5 alignment.

**Additional File 5:** References of previously characterized FLS and DFR included in phylogenetic tree analyses.

**Additional File 6:** Examination of codons at position 133 within the DFR_A_ coding sequence in *Fragaria vesca* and *Fragaria ananassa*, along with a comparison against closely related DFR_N_ sequences. The codons and corresponding amino acids at position 133 are highlighted in red. The color-coded scheme represents different residues: Asparagine (light purple), Alanine (deep purple), and other amino acids (gray) at position 133.

**Additional File 7:** Phylogenetic trees of FLS. (1) FLS tree showing outgroups constructed by IQ-TREE based on an MAFFT alignment, (2a-f) FLS-specific trees with collapsed outgroups. Gymnosperm, monocot, and dicot species are denoted by light blue, light pink, and light green color stripes, respectively. Non-FLS sequences are represented by dashed gray branches while the functional FLSs are indicated by solid black branches. The background color highlights different presumably substrate-preference-determining amino acid residues at position 132: histidine (pale yellow), phenylalanine (dark orange), and tyrosine (dark golden). The hypothesized preferred substrate of the FLS type is written in brackets, DHK, dihydrokaempferol; DHQ, dihydroquercetin and DHM, dihydromyricetin. Distinct clusters of FLSs from major plant orders are labeled for reference. FLS sequences identified in previous studies are highlighted by an asterisk at the start of the terminal branch, with asterisks of functional FLS genes colored in red. (2a) Constructed by IQ-TREE based on a MAFFT alignment, (2b) constructed by IQ-TREE based on a Muscle5 alignment, (2c) constructed by FastTree2 based on a MAFFT alignment, (2d) constructed by FastTree2 based on a Muscle5 alignment, (2e) constructed by MEGA based on a MAFFT alignment, and (2f) constructed by MEGA based on a Muscle5 alignment.

**Additional File 8:** Approximation of dihydroflavonol availability based on gene expression of F3H, F3’H, and F3’5’H in different plant species.

**Additional File 9:** Extended heatmap of 43 species depicting the transcript abundance of analyzed genes in different tissues. DFR* refers to DFR sequences where the residue 133 is not N, D, or A. In *Musa acuminata* C133 residue, *Lotus japonicus* S133, and *Carica papaya* S133.

