## Supplementary figures and images for "Conserved amino acid residues and gene expression patterns associated with the substrate preferences of the competing enzymes FLS and DFR"

### AdditionalFile4

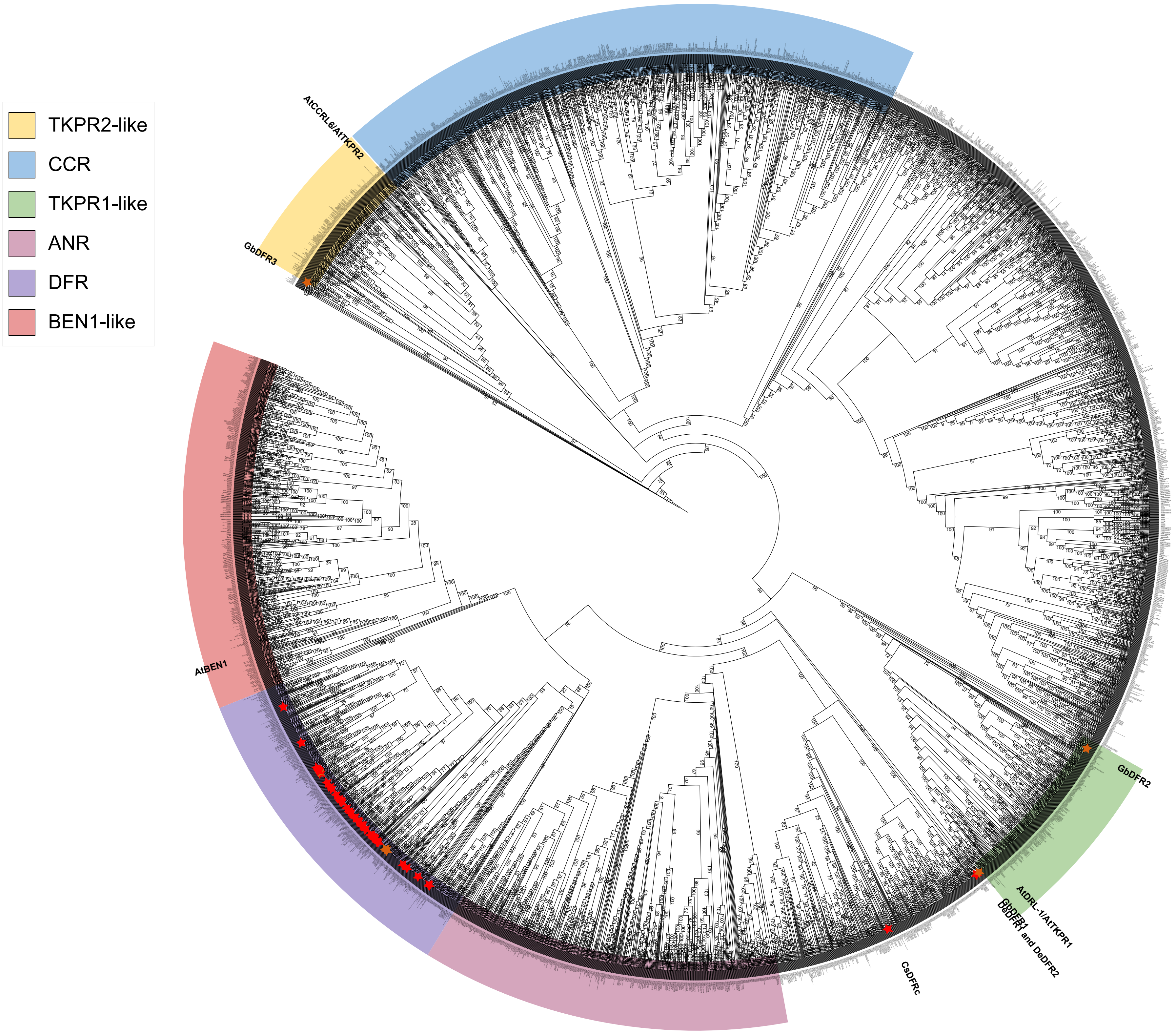

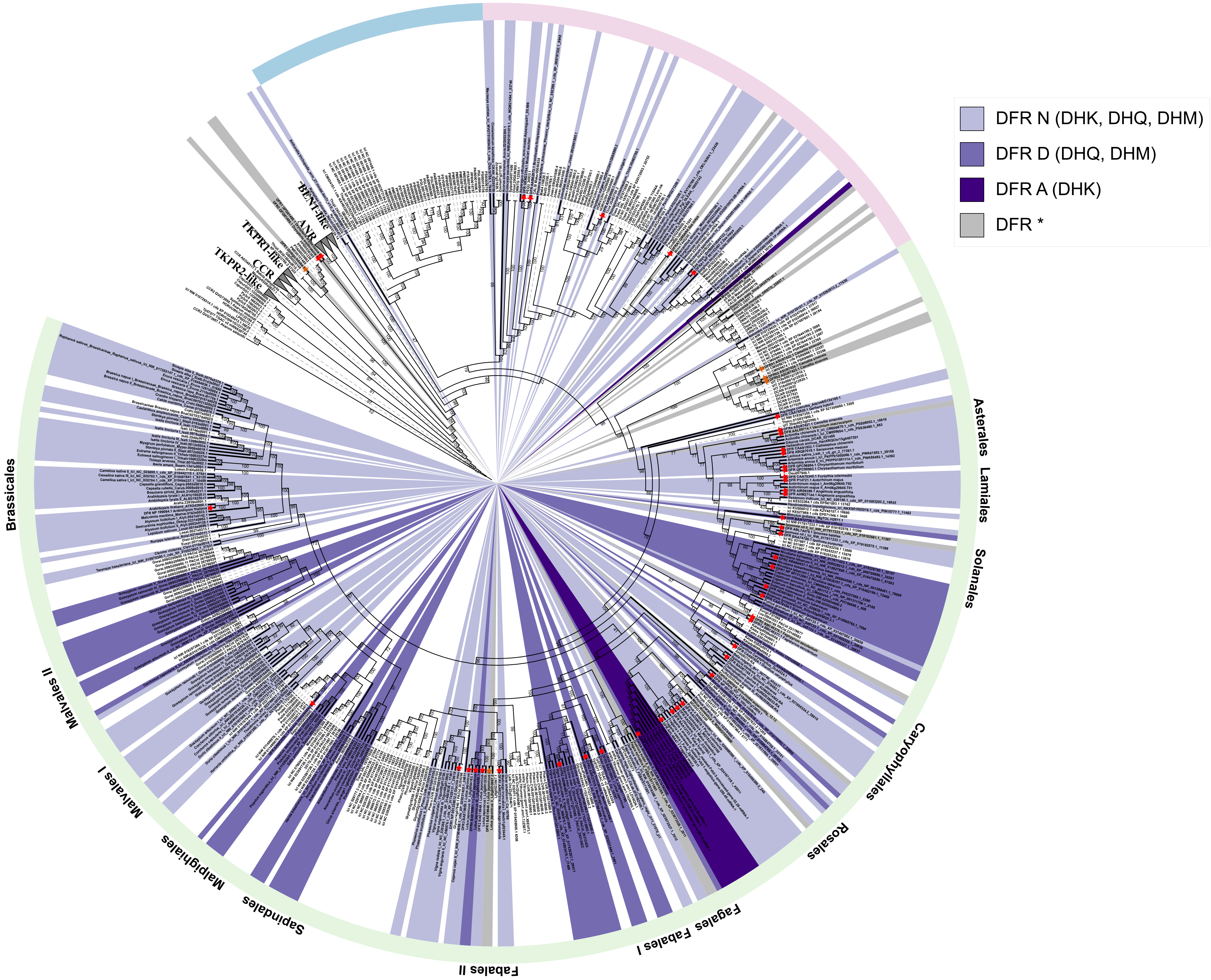

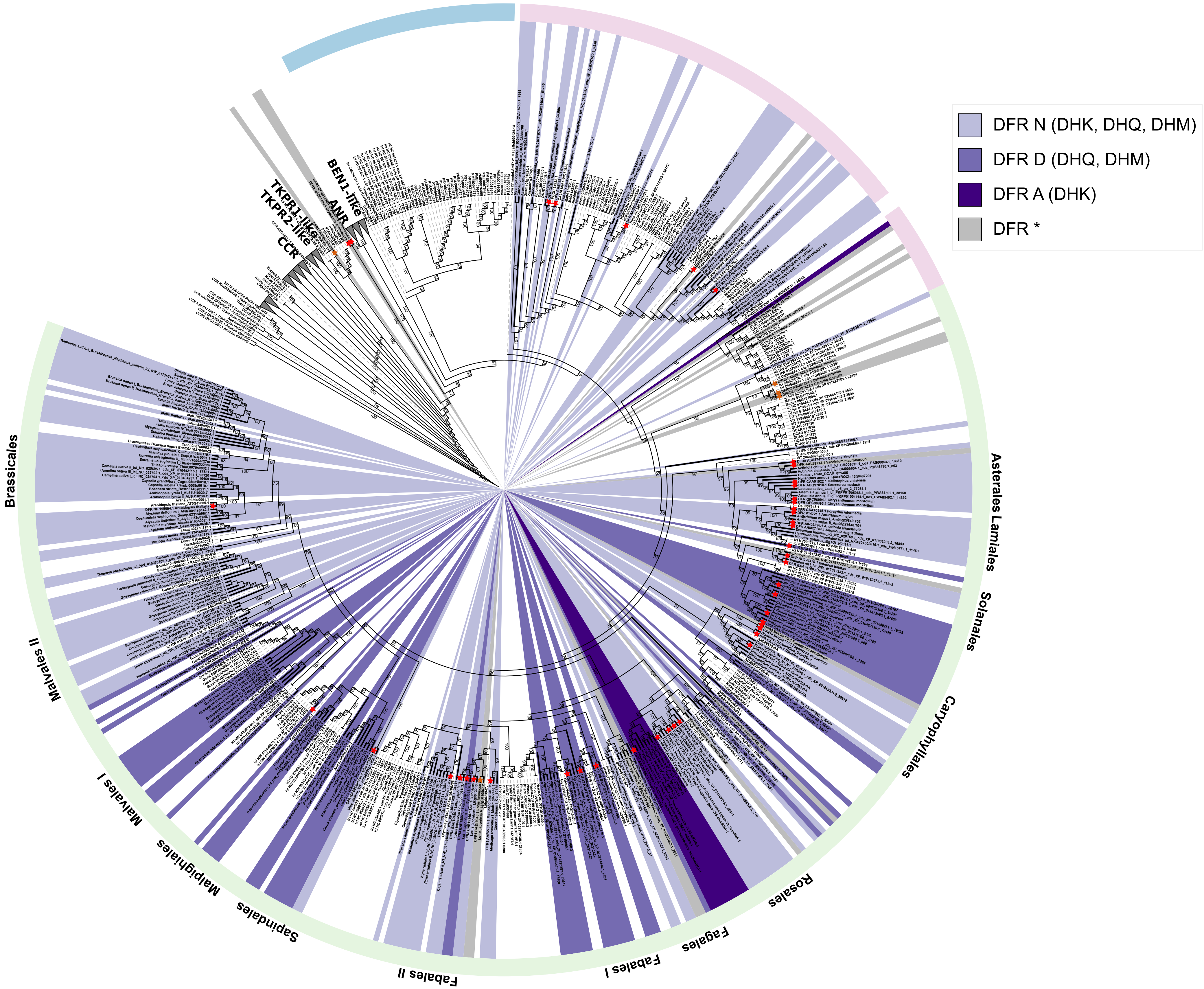

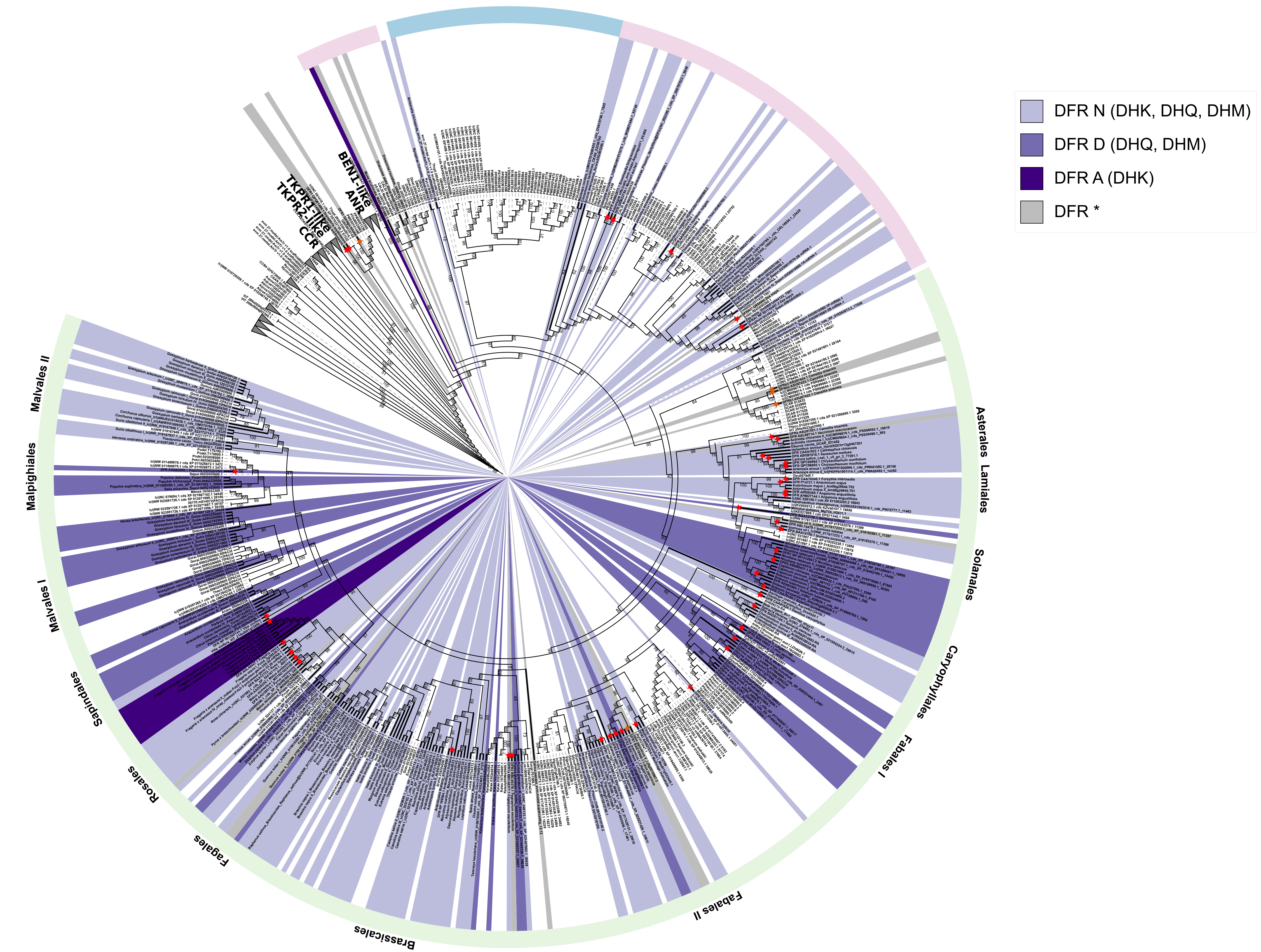

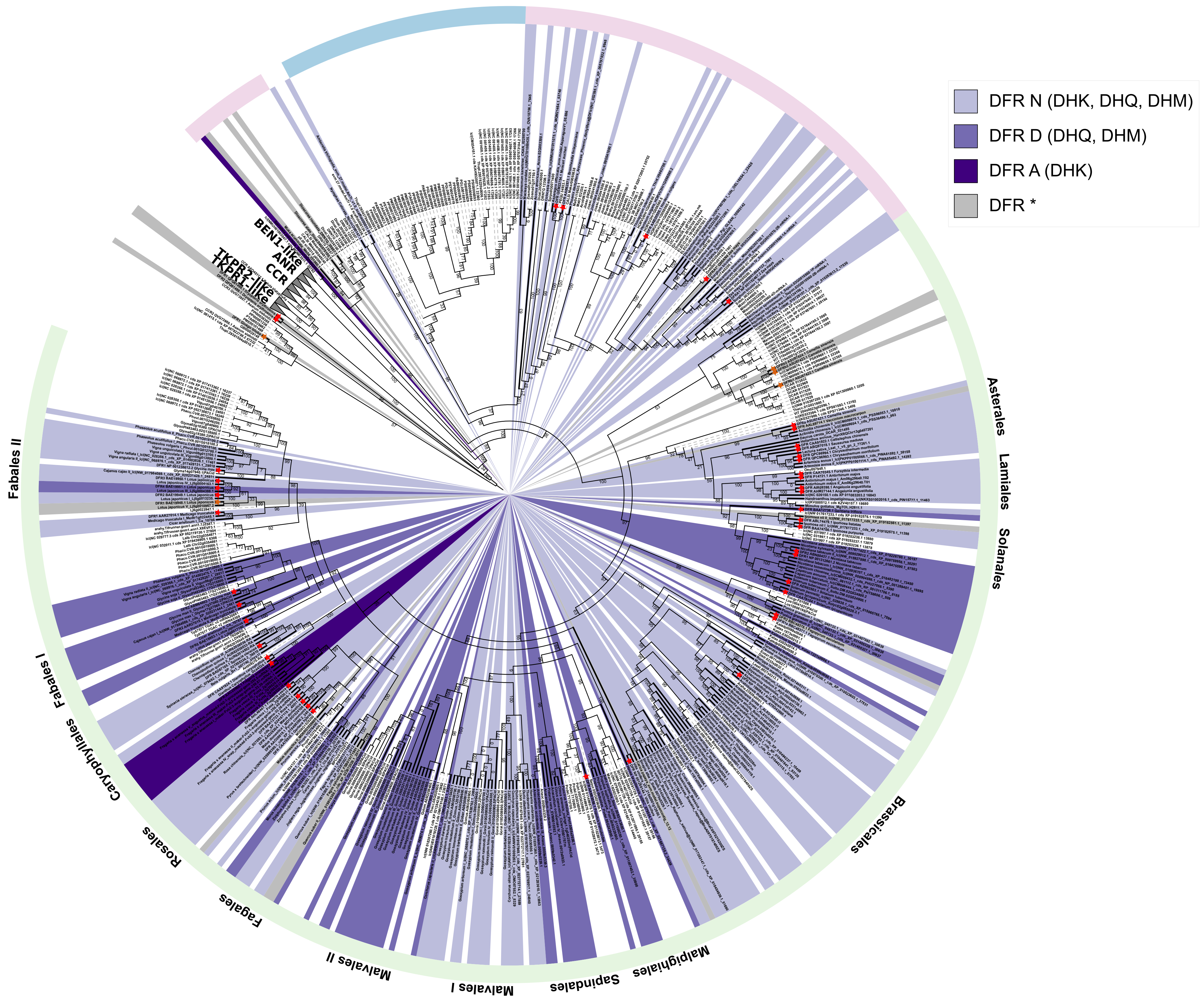

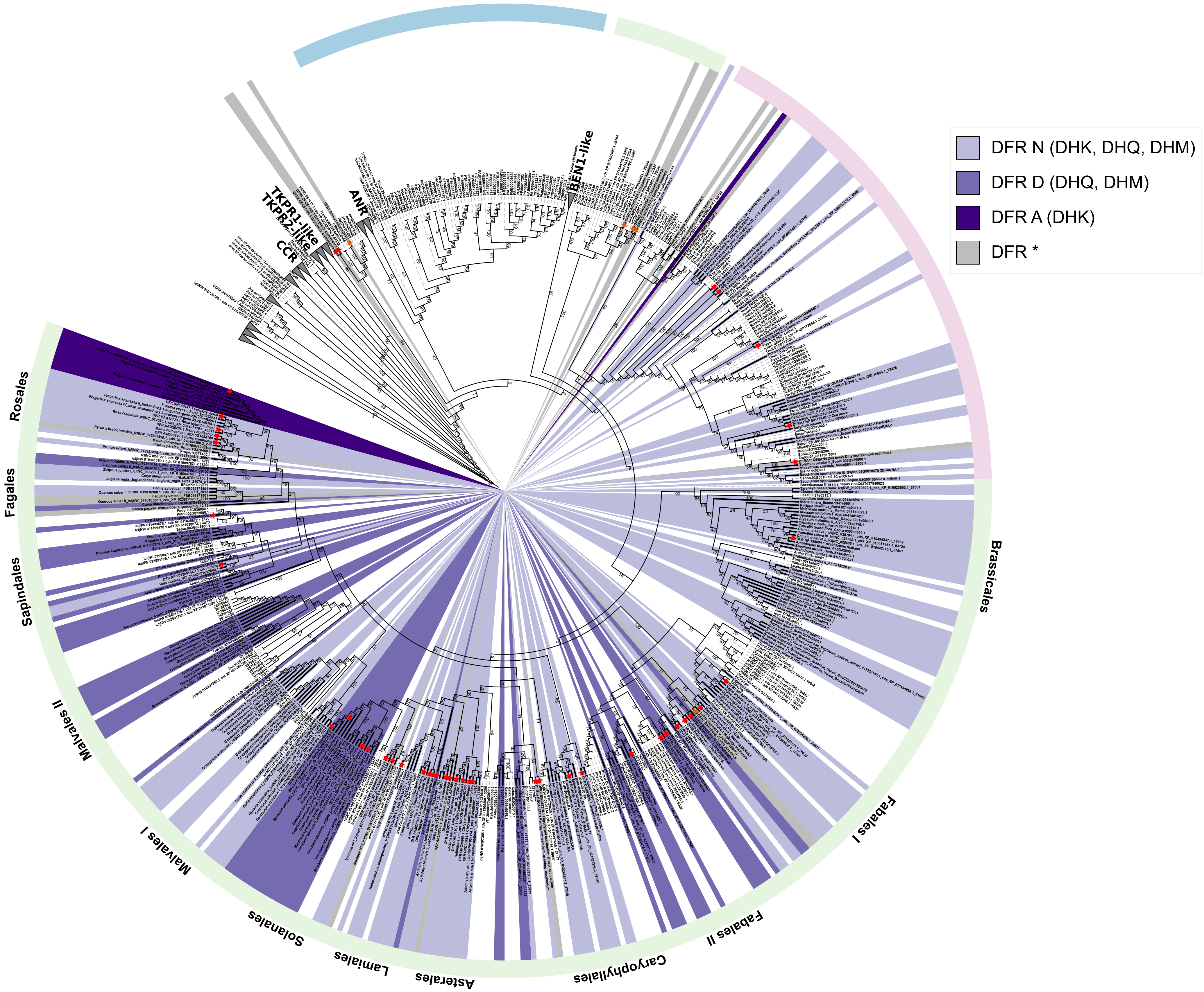

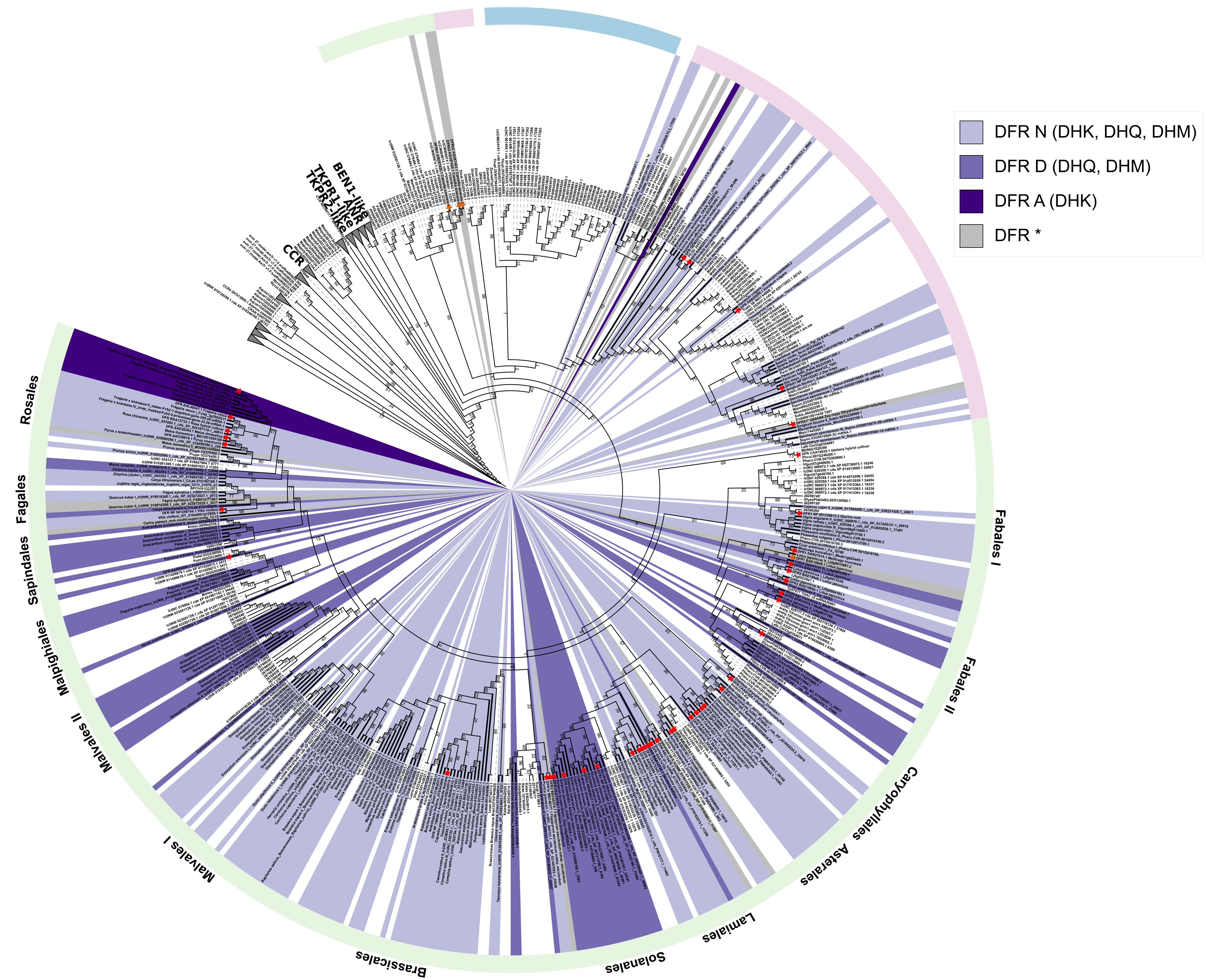

### AdditionalFile6

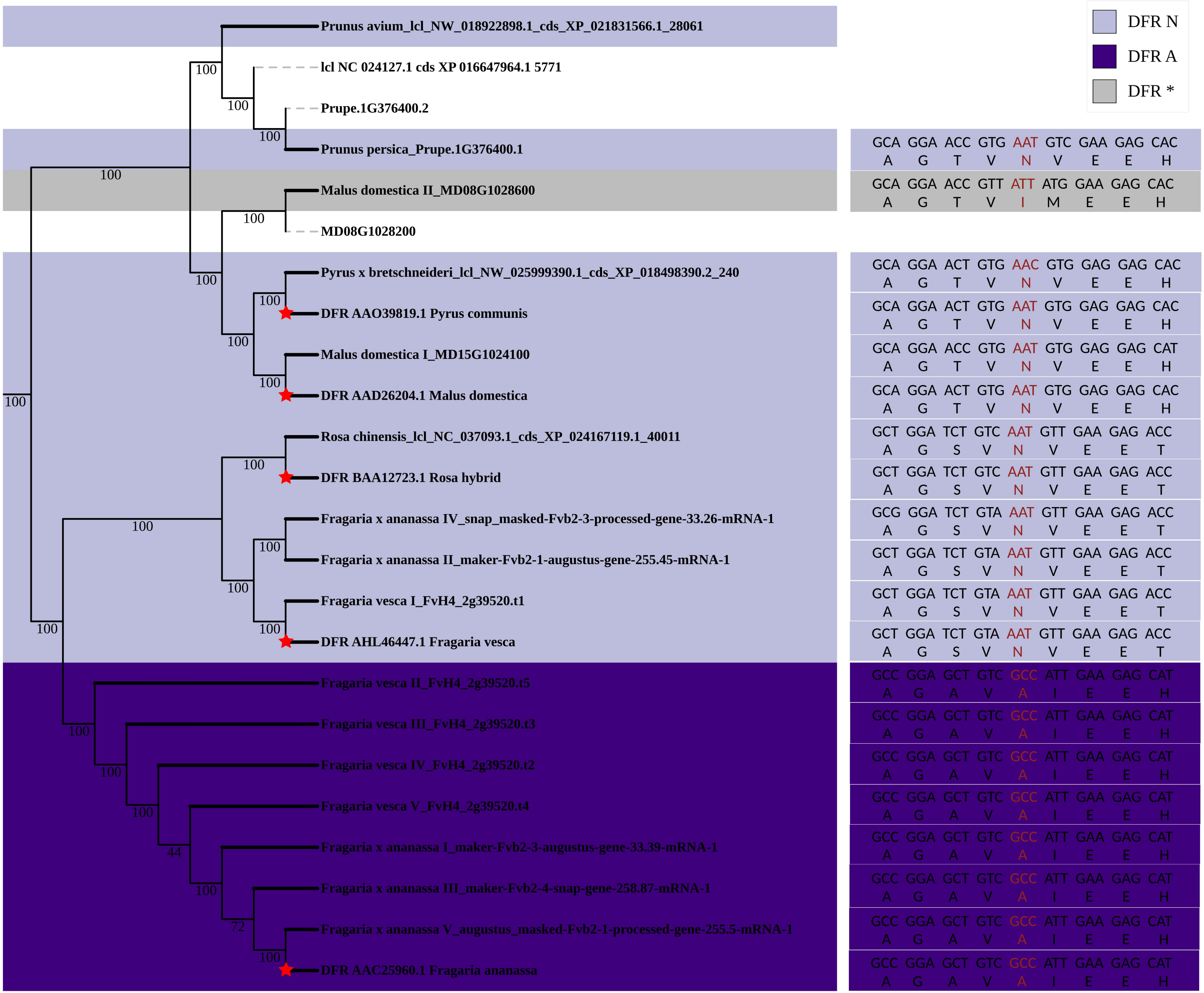

### AdditionalFile7

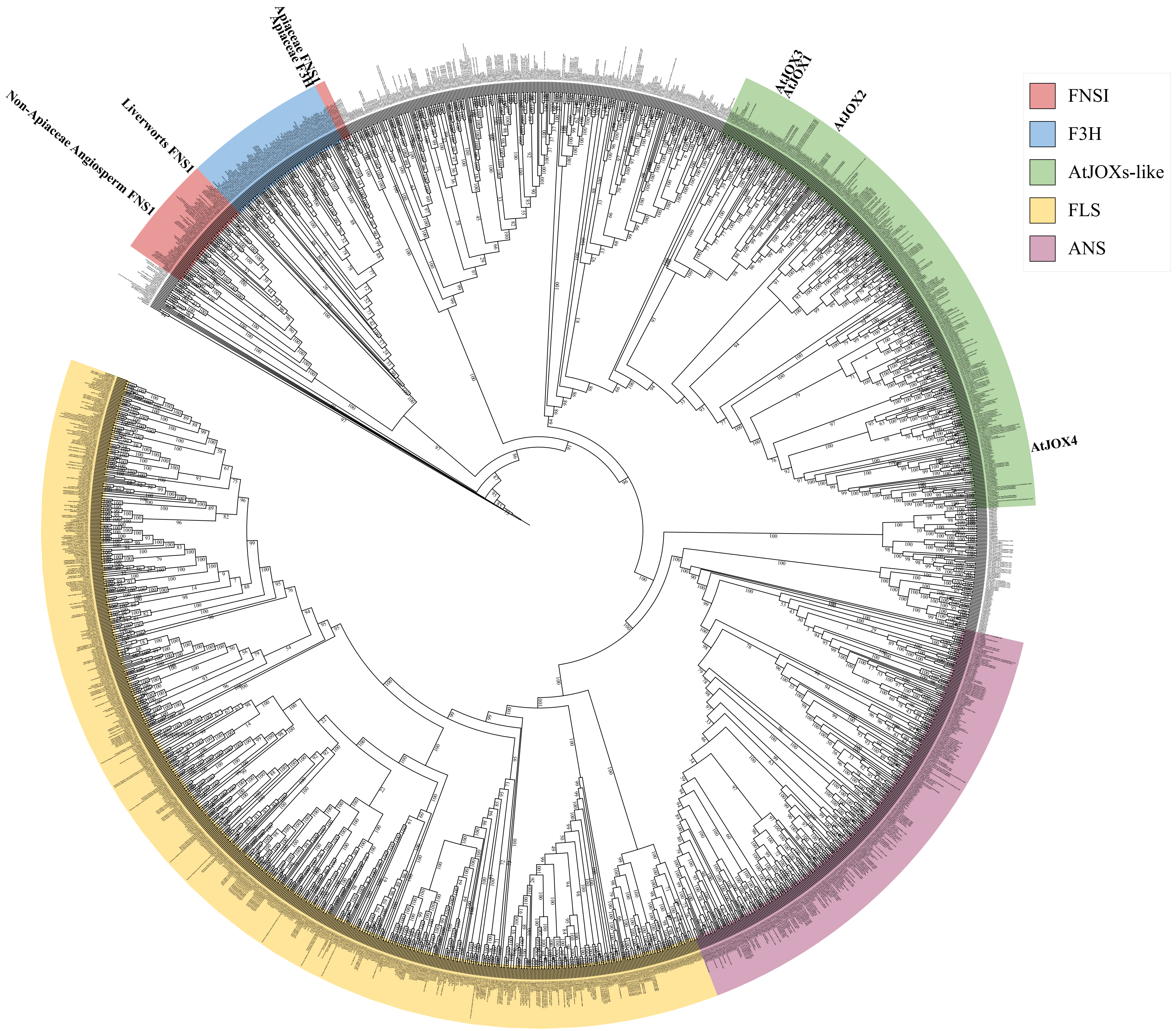

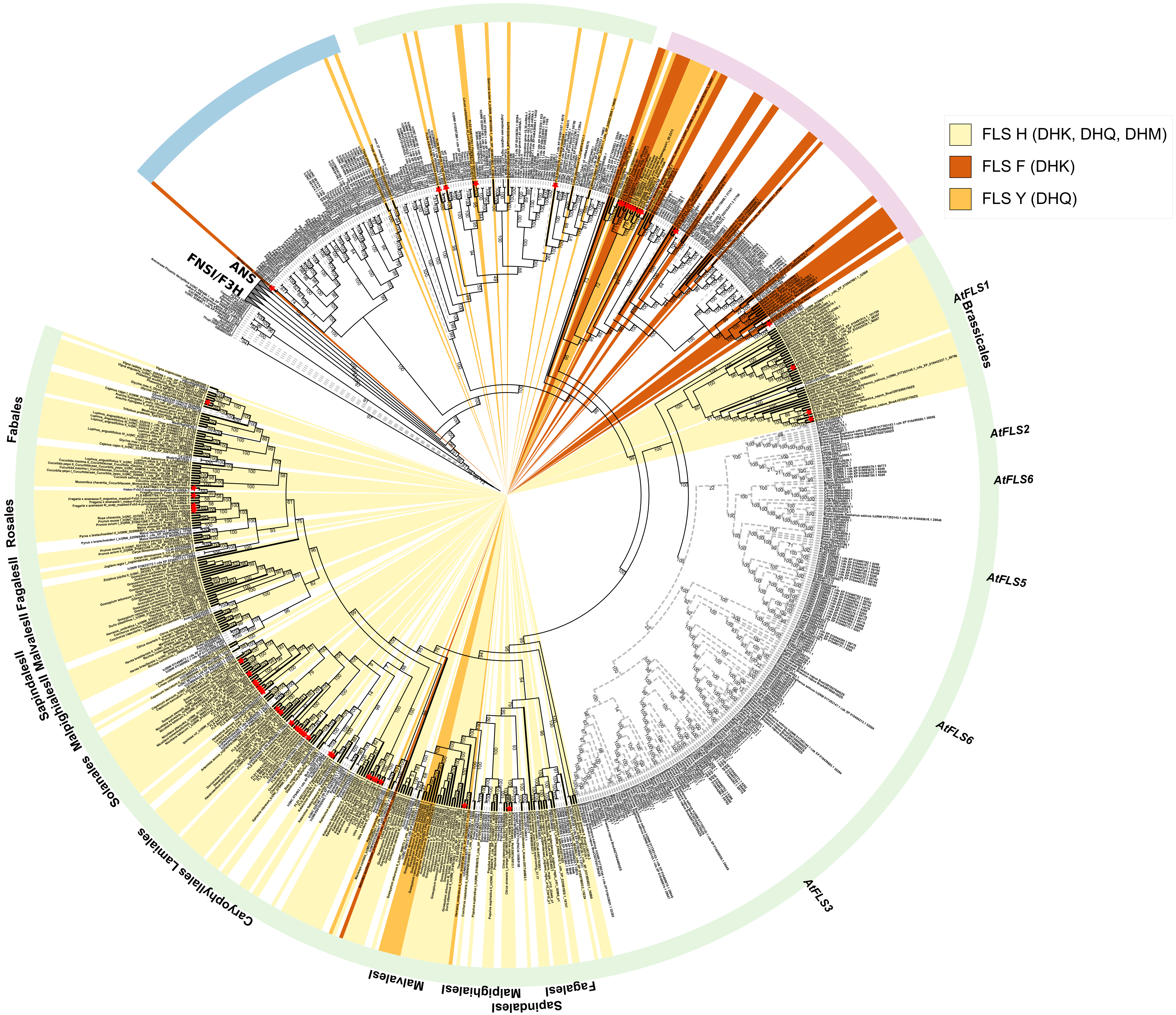

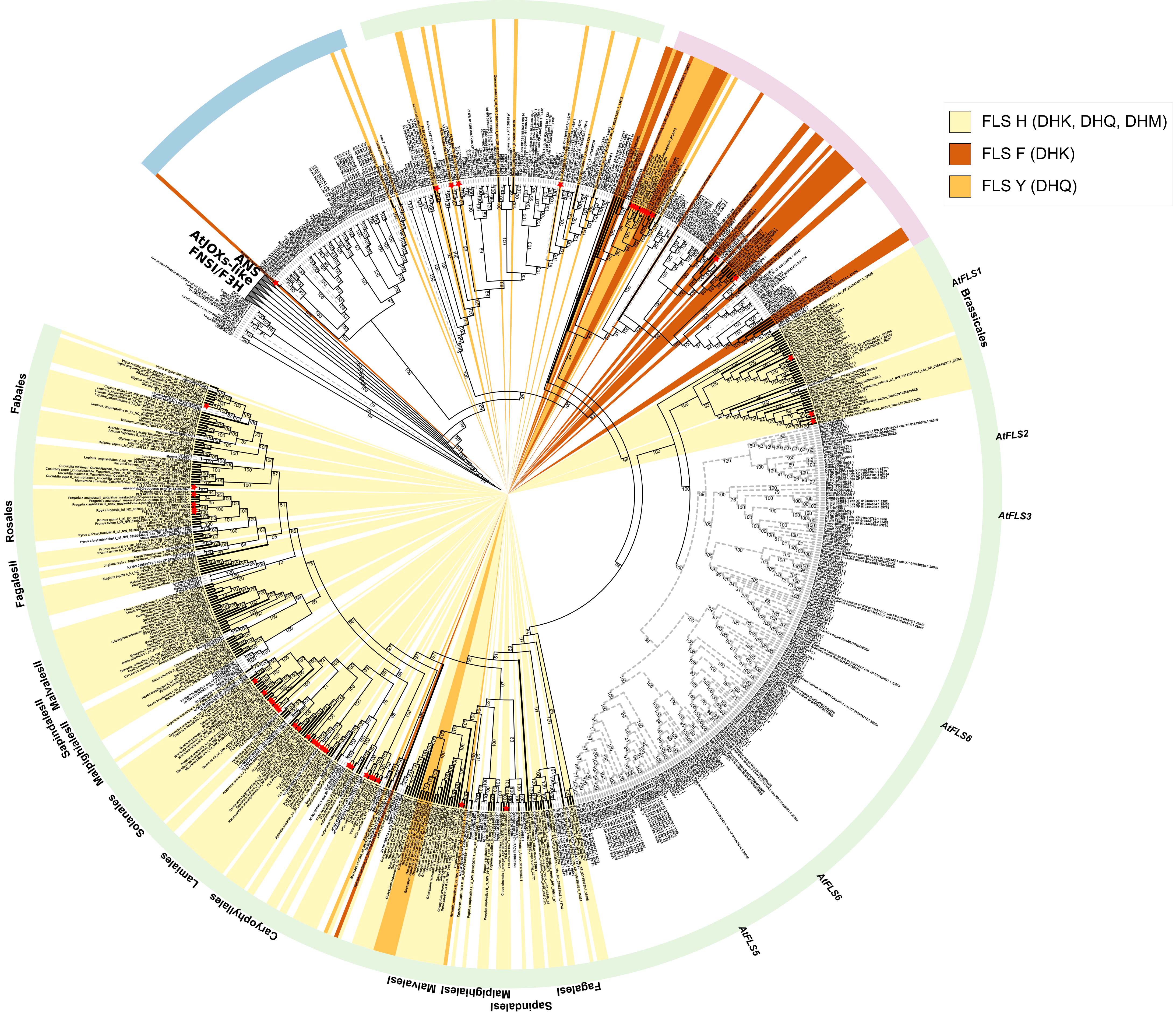

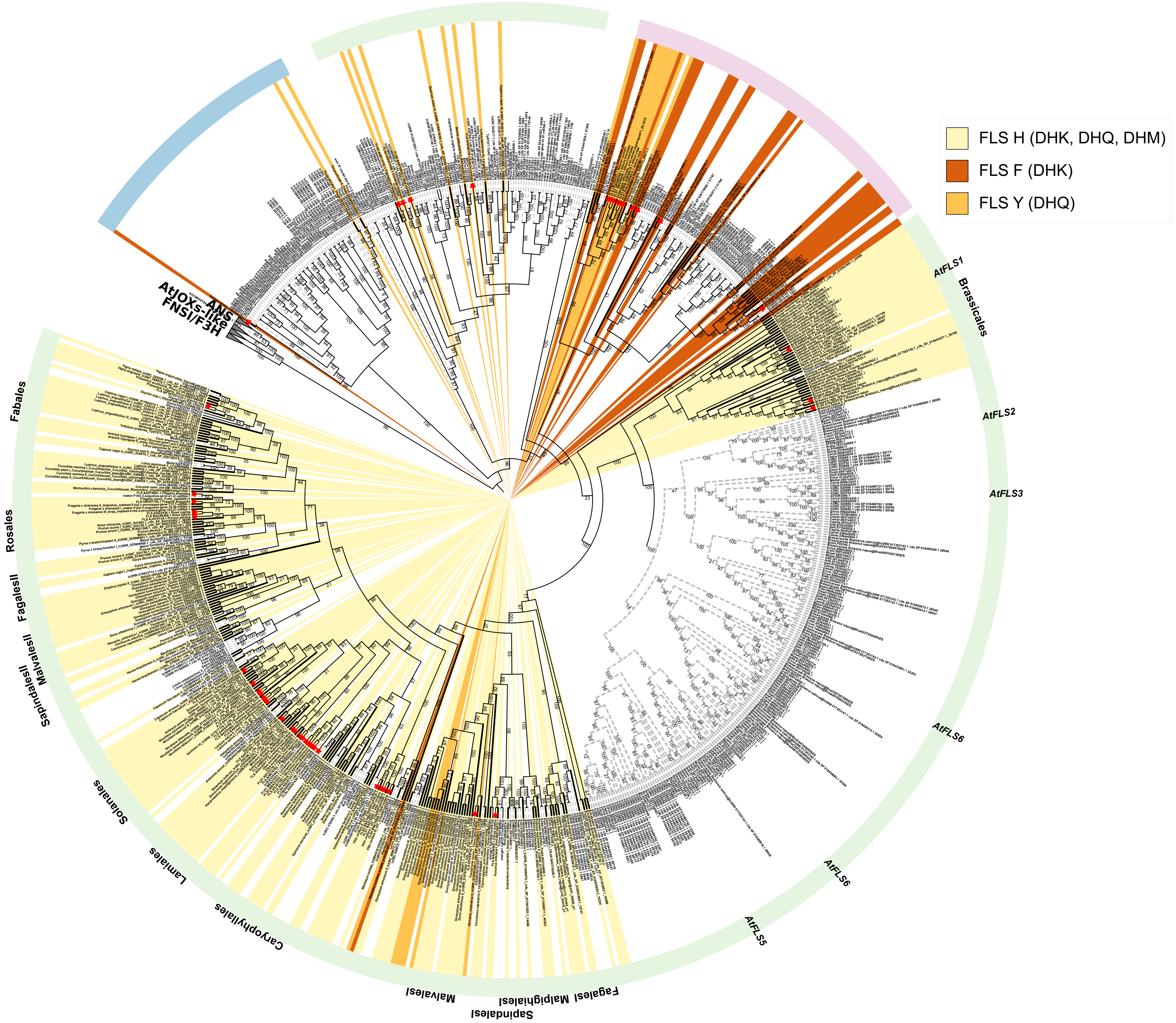

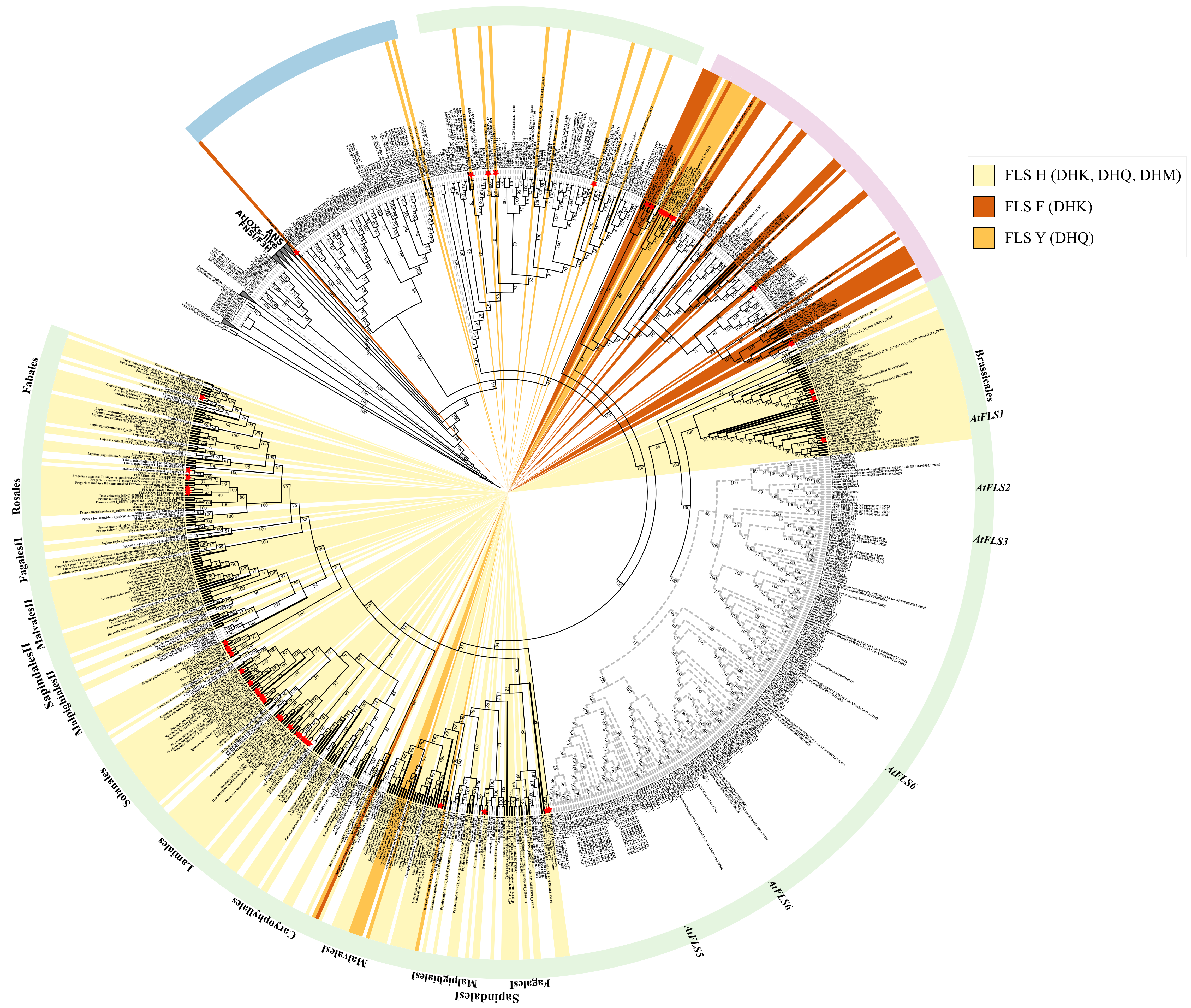

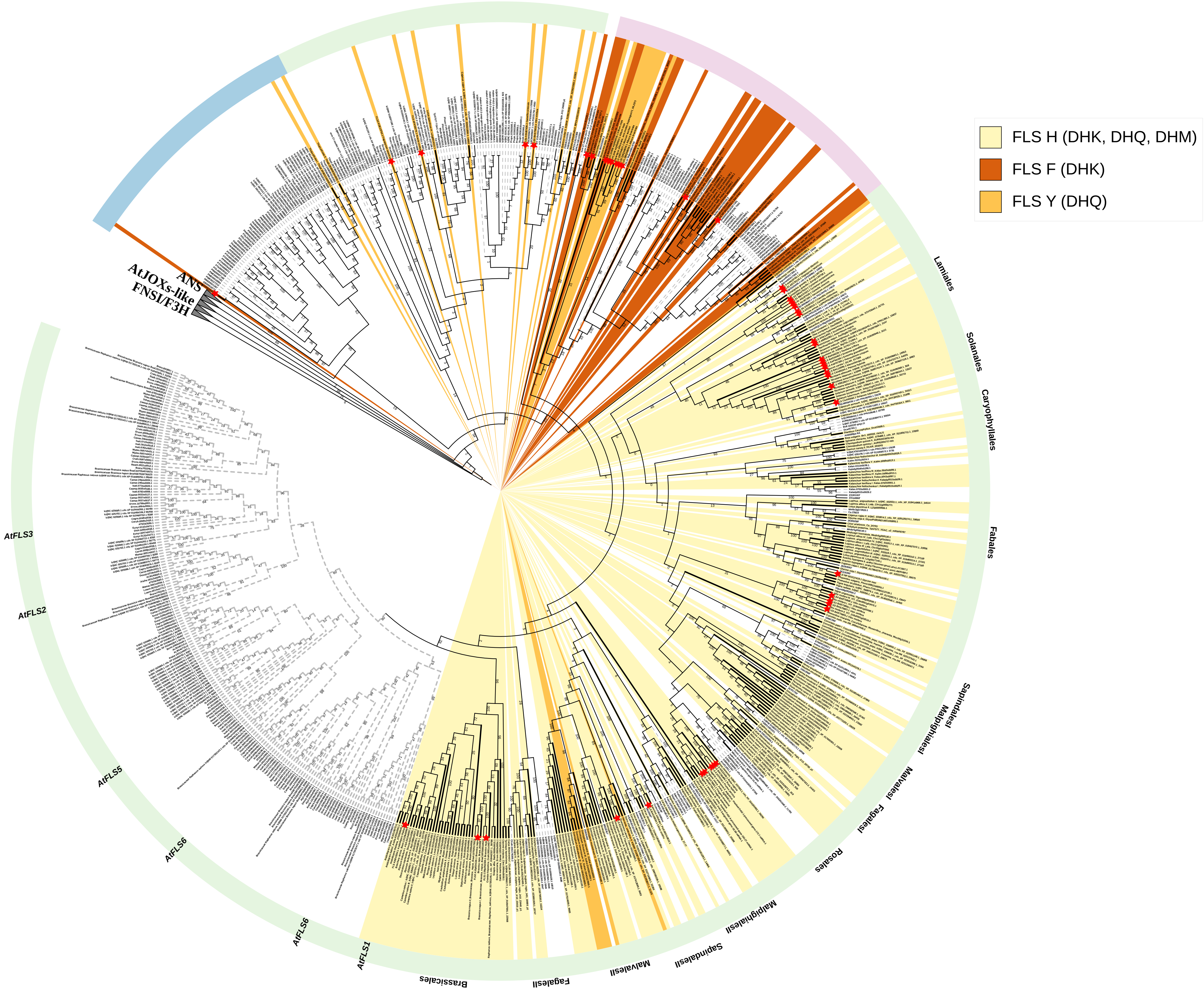

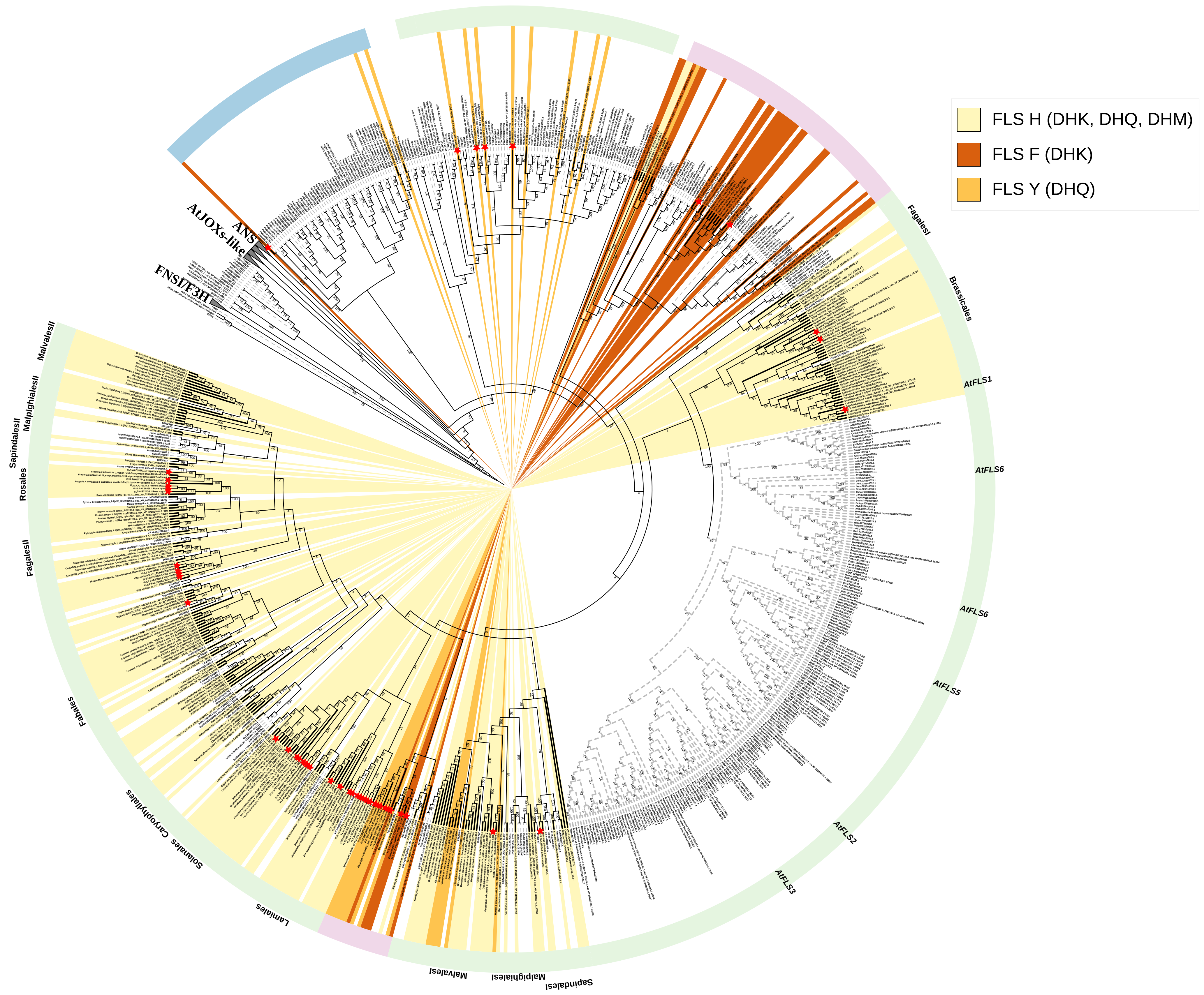

### AdditionalFile9

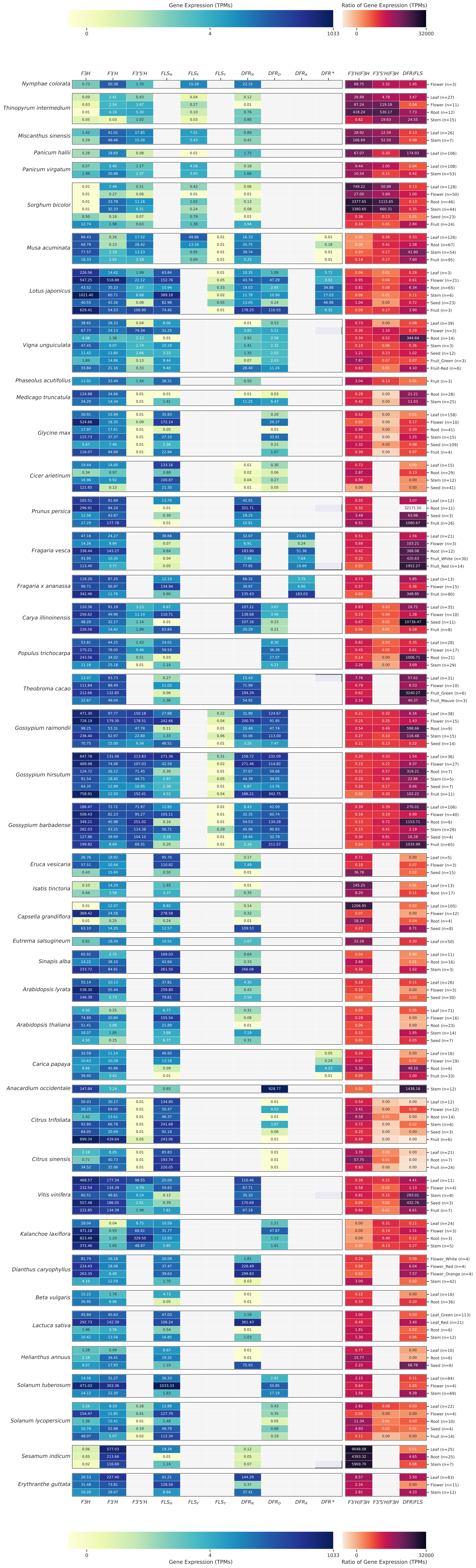
