## AdditionalFile8 for "Conserved amino acid residues and gene expression patterns associated with the substrate preferences of the competing enzymes FLS and DFR"

### Inferring dihydroflavonol availability from relative Flavanone 3-hydroxylase (*F3H)*, Flavonoid 3'-hydroxylase (*F3'H)*, and Flavonoid 3',5'-hydroxylase (*F3'5'H)* gene expression

Major genes like F3H, F3'H, and F3'5'H regulate the branching of flavonol and anthocyanidin biosynthesis by determining the hydroxylation patterns of dihydroflavonols. These genes influence the availability of substrates (DHK, DHQ, DHM) for FLS and DFR enzymes. F3H catalyzes the formation of dihydrokaempferol (DHK), while F3'H adds a hydroxy group converting DHK into dihydroquercetin (DHQ). F3'5'H catalyzes the formation of triple hydroxylated dihydromyricetin (DHM) from DHK/DHQ (**Fig 1**). Understanding the substrate preference of FLS and DFR for differently hydroxylated substrates requires information about the relative availability of DHK, DHQ, and DHM. Since metabolic datasets are not available at the necessary scale, we utilized gene expression data as a semi-quantitative proxy for metabolomics. Transcript abundances are considered as well correlated with the final amount of the gene product because the flavonoid biosynthesis is mostly regulated at the transcriptional level. Therefore, we consider the transcript abundance measured by RNA-seq as a proxy for gene expression. The relative gene expression of *F3H*, *F3'H*, and *F3'5'H* is analyzed to infer the relative production of the three flavonol/anthocyanin precursors, i.e., DHK, DHQ, and DHM. The ratio of *F3'H* to *F3H* and *F3'5'H* to *F3H* were calculated across 43 plant species. For example, a high *F3'H* to *F3H* ratio would suggest that DHQ accounts for most dihydroflavonols. These gene expression ratios were connected to the expression levels and substrate-preference determining amino acid residues in the proteins encoded by *FLS* and *DFR*.

To investigate the gene expression patterns, a heatmap showing data of the above-mentioned genes in 43 species representing the taxonomic diversity of angiosperms was generated (Additional File 9). In **Supplementary Fig 1**, only 10 species are presented to reduce the complexity. The heatmap covers several tissue samples such as leaf, root, stem, flower, fruit, and seed. The samples displayed per individual species depend on data availability. The heatmap analysis revealed distinct expression patterns of *FLS* and *DFR* in different plants. In **Supplementary Fig 1**, the monocot, *Miscanthus sinensis*, demonstrates a higher *F3'H* to *F3H* and *DFR* to *FLS* ratio, suggesting a potential predominance of dihydroquercetin utilization by DFR for anthocyanin or proanthocyanidin production. Cotton exhibits multiple FLS and DFR candidates of $FLS_H$, $FLS_Y$, $DFR_N$, and $DFR_D$ type. Its $FLS_Y$, limited to the Malvales order in the dicot clade, has low gene expression. *Arabidopsis thaliana* lacks *F3'5'H* and exhibits relatively low *DFR* gene expression. The *F3H* and *FLS* gene expression follow the same patterns, suggesting a preference for DHK as a substrate. Species with exclusive $DFR_D$ (aspartate), like oranges and tomatoes, exhibit low *DFR* gene expression despite significant *F3'H* gene expression. In *Dianthus caryophyllus*, red and orange flowers exhibit significant upregulation in *F3H* and $DFR_N$ expression compared to the white flower, while *FLS* expression remains relatively constant. Interestingly, F3'H expression is the least in orange flowers suggesting that its $DFR_N$ is predominantly acting on DHK as substrate leading to the formation of pelargonidin-based anthocyanins, hence the orange colour. The red flower has high gene expression of both F3H and F3'H suggesting $DFR_N$ here might be catalyzing both DHK and DHQ, leading to the production of cyanidin- and pelargonidin-based pigments and contributing to the observed red phenotype.

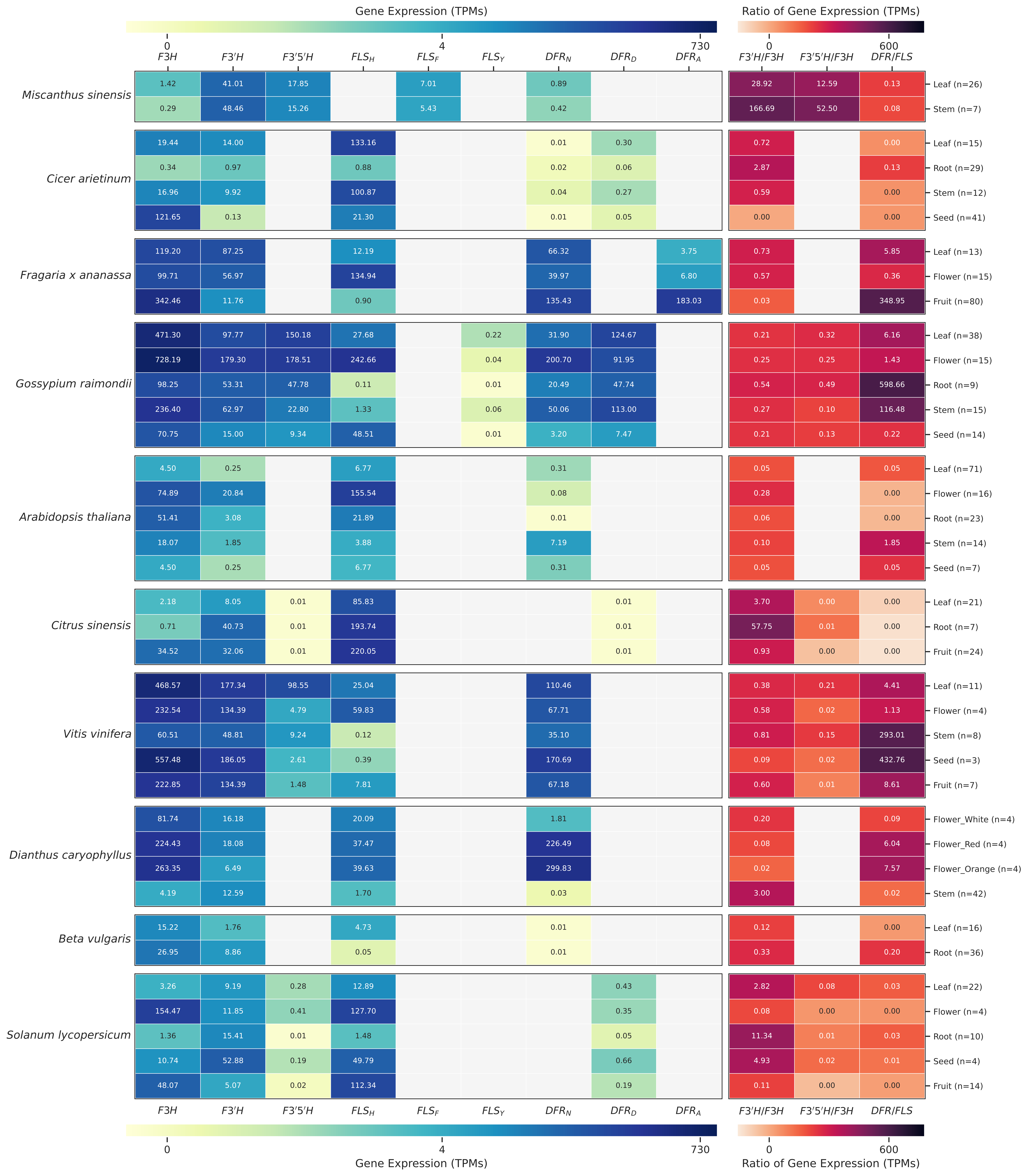

Gene Expression (TPMs)
0
4
730
Ratio of Gene Expression (TPMs)
0
600
F3H
F3′H
F3′5′H
FLS_H
FLS_F
FLS_Y
DFR_N
DFR_D
DFR_A
F3′H/F3H
F3′5′H/F3H
DFR/FLS
Miscanthus sinensis
1.42 41.01 17.85 7.01 0.89 28.92 12.59 0.13 Leaf (n=26)
0.29 48.46 15.26 5.43 0.42 166.69 52.50 0.08 Stem (n=7)
Cicer arietinum
19.44 14.00 133.16 0.01 0.30 0.72 0.00 Leaf (n=15)
0.34 0.97 0.88 0.02 0.06 2.87 0.13 Root (n=29)
16.96 9.92 100.87 0.04 0.27 0.59 0.00 Stem (n=12)
121.65 0.13 21.30 0.01 0.05 0.00 0.00 Seed (n=41)
Fragaria x ananassa
119.20 87.25 12.19 66.32 3.75 0.73 5.85 Leaf (n=13)
99.71 56.97 134.94 39.97 6.80 0.57 0.36 Flower (n=15)
342.46 11.76 0.90 135.43 183.03 0.03 348.95 Fruit (n=80)
Gossypium raimondii
471.30 97.77 150.18 27.68 0.22 31.90 124.67 0.21 0.32 6.16 Leaf (n=38)
728.19 179.30 178.51 242.66 0.04 200.70 91.95 0.25 0.25 1.43 Flower (n=15)
98.25 53.31 47.78 0.11 0.01 20.49 47.74 0.54 0.49 598.66 Root (n=9)
236.40 62.97 22.80 1.33 0.06 50.06 113.00 0.27 0.10 116.48 Stem (n=15)
70.75 15.00 9.34 48.51 0.01 3.20 7.47 0.21 0.13 0.22 Seed (n=14)
Arabidopsis thaliana
4.50 0.25 6.77 0.31 0.05 0.05 Leaf (n=71)
74.89 20.84 155.54 0.08 0.28 0.00 Flower (n=16)
51.41 3.08 21.89 0.01 0.06 0.00 Root (n=23)
18.07 1.85 3.88 7.19 0.10 1.85 Stem (n=14)
4.50 0.25 6.77 0.31 0.05 0.05 Seed (n=7)
Citrus sinensis
2.18 8.05 0.01 85.83 0.01 3.70 0.00 0.00 Leaf (n=21)
0.71 40.73 0.01 193.74 0.01 57.75 0.01 0.00 Root (n=7)
34.52 32.06 0.01 220.05 0.01 0.93 0.00 0.00 Fruit (n=24)
Vitis vinifera
468.57 177.34 98.55 25.04 110.46 0.38 0.21 4.41 Leaf (n=11)
232.54 134.39 4.79 59.83 67.71 0.58 0.02 1.13 Flower (n=4)
60.51 48.81 9.24 0.12 35.10 0.81 0.15 293.01 Stem (n=8)
557.48 186.05 2.61 0.39 170.69 0.09 0.02 432.76 Seed (n=3)
222.85 134.39 1.48 7.81 67.18 0.60 0.01 8.61 Fruit (n=7)
Dianthus caryophyllus
81.74 16.18 20.09 1.81 0.20 0.09 Flower_White (n=4)
224.43 18.08 37.47 226.49 0.08 6.04 Flower_Red (n=4)
263.35 6.49 39.63 299.83 0.02 7.57 Flower_Orange (n=4)
4.19 12.59 1.70 0.03 3.00 0.02 Stem (n=42)
Beta vulgaris
15.22 1.76 4.73 0.01 0.12 0.00 Leaf (n=16)
26.95 8.86 0.05 0.01 0.33 0.20 Root (n=36)
Solanum lycopersicum
3.26 9.19 0.28 12.89 0.43 2.82 0.08 0.03 Leaf (n=22)
154.47 11.85 0.41 127.70 0.35 0.08 0.00 0.00 Flower (n=4)
1.36 15.41 0.01 1.48 0.05 11.34 0.01 0.03 Root (n=10)
10.74 52.88 0.19 49.79 0.66 4.93 0.02 0.01 Seed (n=4)
48.07 5.07 0.02 112.34 0.19 0.11 0.00 0.00 Fruit (n=14)

**Supplementary Fig 1:** Gene expression heatmap and gene expression ratio analysis of 10 selected species across various tissue samples. White tiles indicate that no gene was discovered for a certain function/type. The heatmap tiles represent the actual expression values in TPMs, with the number of samples indicated on the right. The expression heatmap shows genes associated with the branching of *FLS* and *DFR* within the flavonoid biosynthesis. Blue and yellow colors indicate high and low expression levels, respectively (see color bar). On the right, a heatmap illustrates the ratios of gene expression to investigate the substrate hydroxylation pattern in the FLS-DFR branching. Dark and light colors indicate high and low expression levels, respectively, based on the color bar. A full heatmap with all 43 investigated species is available in Additional File 8.

From the gene expression analysis (Additional File 9), we distilled five important observations. Firstly, we observed diverse substrate preferences among multiple DFR/FLS copies, as seen in *Lotus japonicus* and various *Gossypium* species harboring both $DFR_N$ and $DFR_D$, as well as $FLS_H$ and $FLS_Y$, with $DFR_N$ favoring DHK and $DFR_D$ preferring DHQ. The $FLS_Y$ in *L. japonicus* is an ancestral FLS (aFLS) while the ones in *Gossypium* are re-emerged $FLS_{Y.}$ Interestingly, $FLS_Y$ in both species exhibited almost negligible expression. Secondly, tissue-specific expression of *FLS* and *DFR* was evident, as seen in *Fragaria ananassa*, where *FLS* displayed high expression in white flowers while *DFR* was more expressed in red fruits. This tissue-specific expression of *FLS* and *DFR* would mitigate competition between both in the same tissue. Thirdly, a preference for flavonols or anthocyanins might also affect the metabolic flux. Legume species like *Vigna unguiculata*, *Phaseolus acutifolius*, *Glycine max*, and *Cicer arietinum* from Fabales orders are known to produce flavonoids and isoflavonoids since they have a physiological role in nodulation [1]. They were found to have a greater gene expression ratio of *FLS* to *DFR*, while others like *Vitis vinifera* with a high anthocyanin content [2] have a higher *DFR* to *FLS* expression ratio. Fourthly, the presence of an alternate pigmentation pathway may also influence the *DFR* expression levels. The betalain-pigmented *Beta vulgaris* does not accumulate anthocyanins [3] and hence, shows no *DFR* expression. Similar patterns were observed in some high-carotenoid species like *Solanum lycopersicum* and *Carica papaya*, which exhibited minimal to no *DFR* gene expression. Lastly, we observed that the expression of the same *FLS*/*DFR* type in the same plant tissue with different phenotypes differed. For example, *Vigna unguiculata* green fruit showed lower expression of $DFR_N$, $DFR_D$, *F3'H,* and *F3H* compared to red fruit, indicating increased substrate availability in red fruit. Despite this, *FLS* expression remained constant in both red and green fruit. This pattern was consistent in other plants, in the white and red fruit of *Fragaria vesca*, green and red leaf of *Lactuca sativa,* and white, red, and orange flowers of *Dianthus caryophyllus*, suggesting a complex regulation mechanism. Previous studies suggested feedback mechanisms in flavonoid biosynthesis regulation [4–6]. Another possible explanation could be the influence of anthocyanin biosynthesis transcription factors like the MBW complex [7]. Surprisingly, in *Theobroma cacao*, green fruit exhibited higher expression of *F3H*, *F3'H*, and $DFR_N$ compared to mauve fruit. This apparent disparity in the gene expression and observed phenotype may be explained by a required expression of the anthocyanin biosynthesis genes before the anthocyanin pigmentation can become visible. Previous studies have reported that the accumulation of flavonols/anthocyanins may not always be directly correlated with the expression levels of *FLS*/*DFR* [8,9]. This can be explained because the accumulation of anthocyanins is preceded by the expression of the required biosynthesis genes [10,11], i.e., enzymes need to be available for anthocyanin production.

### Method

The accession numbers of different tissues (leaf, stem, flower, root, seed, and fruit) for the 43 species were retrieved from the SRA (SRA IDs used in this study available at https://github.com/bpucker/DFR_vs_FLS). This sample selection was further reduced to avoid the inclusion of any specific stress treatments, i.e., the different samples should represent different plant parts under 'normal' conditions. We aimed to assess the expression of a specific gene in a particular sample. The Python script generate_subset.py [12] was deployed to extract an expression data subset from the count table containing exclusively the values belonging to the accession numbers of the desired samples. The transcripts per million (TPMs) belonging to isoforms of a gene and close paralogs were aggregated per RNA-seq sample as previously described [13]. To obtain an overall representation of the expression of a gene, we calculated the median value of the aggregated TPMs across a group of samples (**Supplementary Fig 2**). The median was chosen due to its tolerance towards outliers. To visualize the gene expression patterns, a heatmap was generated using the seaborn package [14] in Python. The heatmap depicted the relative gene expression levels of *F3H*, *F3'H*, *F3'5'H*, *FLS*, and *DFR*. Additionally, ratios of *F3'H*/*F3H*, *F3'5'H*/*F3H*, and *DFR*/*FLS* were plotted to provide insights into their relative expression levels and potential interdependencies within the flavonoid biosynthesis.

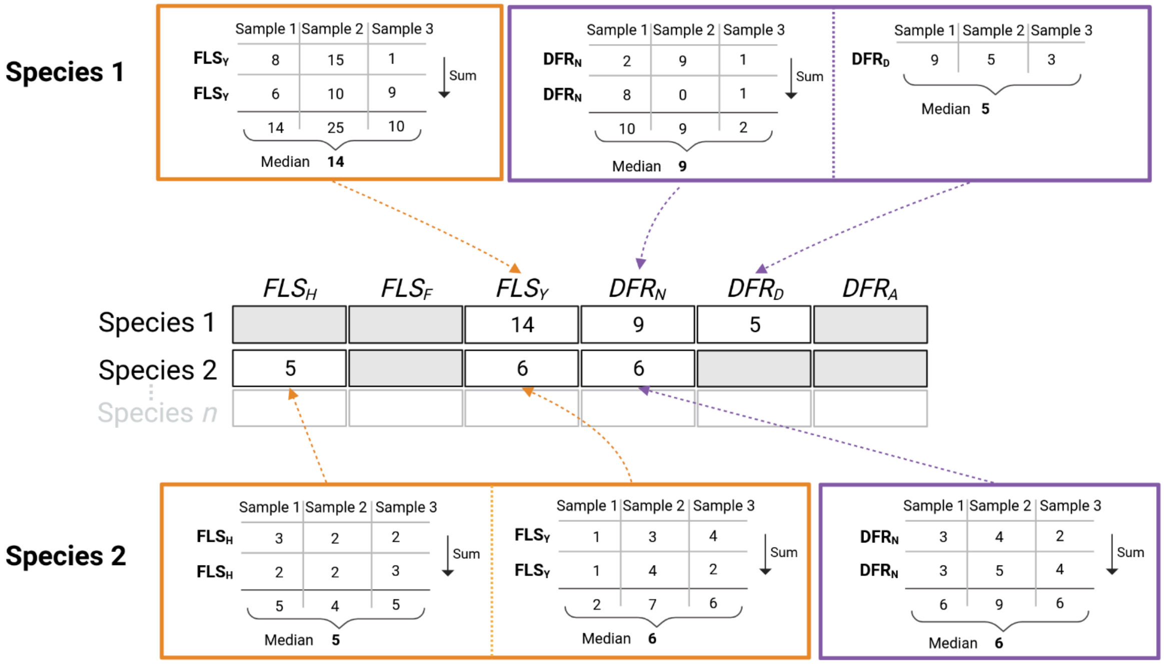


**Supplementary Fig 2:** Illustration of the cross-species gene expression comparison. All displayed values are TPMs. The values of close paralogs were replaced by their sum if these paralogs had the same substrate affinity determining residue at the most important position.
